# Computational exploration of treadmilling and protrusion growth observed in fire ant rafts

**DOI:** 10.1101/2021.01.05.425514

**Authors:** Robert J. Wagner, Franck J. Vernerey

## Abstract

Condensed active matter systems regularly achieve cooperative emergent functions that individual constituents could not accomplish alone. The rafts of fire ants (Solenopsis invicta) are often studied in this context for their ability to create structures comprised entirely of their own bodies, including tether-like protrusions that facilitate exploration of flooded environments. While similar protrusions are observed in cytoskeletons and cellular aggregates, they are generally dependent on morphogens or external gradients leaving the isolated role of local interactions poorly understood. Here we demonstrate through an ant-inspired, agent-based numerical model how protrusions in ant rafts may emerge spontaneously due to local interactions and how phases of exploratory protrusion growth may be induced by increased ant activity. These results provide an example in which functional morphogenesis of condensed active matter may emerge purely from locally-driven collective motion and may provide a source of inspiration for the development of autonomous active matter and swarm robotics.

## Introduction

Condensed active matter is defined as an ensemble of constituents that consume energy to do mechanical work while remaining entangled or in physical contact with one another (Hu et al., 2016). Examples of such systems are omnipresent in nature and include systems ranging from the cytoskeletal walls of single cells (Neuhaus et al., 1983) to clustered aggregations of worms or insects (Adams et al., 2011; Deblais et al., 2020; Peleg et al., 2018). This material class is exemplary for its ability to display complex cooperative behavior and morph despite remaining cohesively unified. Cytoskeletal walls utilize biased, collective restructuring of their actin meshes to facilitate motility (Bugyi and Carlier, 2010), and clusters of cells undergo collective motion through a plethora of mechanisms that drive processes such as healing of wounds or invasion by tumor cells (Méhes and Vicsek, 2014; Trepat et al., 2012). Recently, we discovered and reported similar emergent behavior in 2D rafts comprised of fire ants (*Solenopsis invicta*) (Wagner et al., 2020). During floods, fire ants condense into buoyant rafts made entirely of worker ant bodies, thereby keeping their colonies unified and bolstering chances of survival (Adams et al., 2011; Mlot et al., 2011). In doing so, fire ants create an intriguing example of nonequilibrium, biphasic active matter that consists of a condensed structural network of inter-linked ants floating on the surface of the water, on top of which dispersed surface ants walk as self-propelled particles (SPPs). We demonstrated that these ant rafts, when anchored to a rod stemming from the water’s surface, display the ability to sprout tether-like protrusions that emerge and recess perpetually over the span of hours (Wagner et al., 2020). The sustained emergence of these growths relies on treadmilling dynamics in which the structural network comprising the raft continually contracts, while freely active ants on the surface of the raft deposit into its edges and drive outwards expansion (**Fig. 1**). The population of surface ants that fuels outwards expansion is continually replenished by unbinding of structural ants from the bulk of the raft and their subsequent phase transition into the freely active layer.

**Figure 1.**
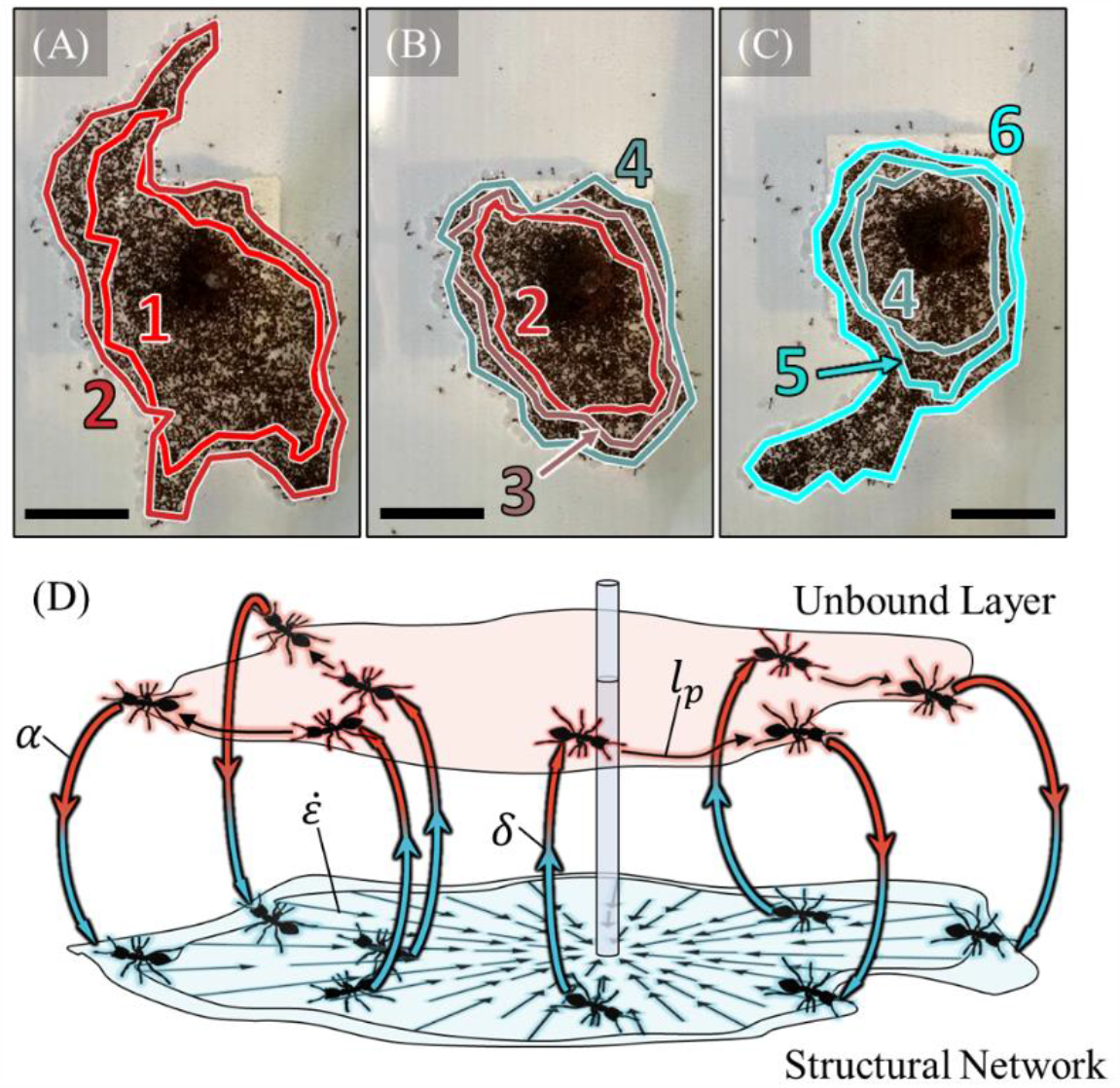
Treadmilling. **(A-C)** A top view of an experimental raft is illustrated at the start **(A)**, middle **(B)** and end **(C)** of a roughly 106 min duration. To visually illustrate treadmilling, the perimeter is traced every 22 minutes by numbered and distinctly colored outlines with 1 representing the oldest set of ants and 6 representing the newest. The shrinking of these contours indicates retraction of the raft structure, while the existence of new layers indicates outwards expansion. Periods of raft expansion and coinciding protrusion emergence **(A**,**C)** were interrupted by interstitial spans of decreased activity and less eccentric morphologies **(B)**. All scale bars represent 10 𝓁 where 𝓁 = 2.93 ± 0.02 mm is the approximate average body length of 1 ant. See **Movie S1** for unannotated video. **(D)** A schematic visually illustrates the four concurrent mechanisms of treadmilling: (1) structural raft contraction at a global rate 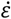, (2) phase transition of structural ants to unbound active ants in the bulk at a nominal rate *δ*, (3) transport of the surface ants on top of the raft with a mean persistence length *l*_*p*_, and (4) binding of surface ants back into the structural network at the edges of the raft at nominal rate *α*. The schematic is taken from Wagner et al. (2020).

On occasion, fire ants utilize protrusions as floating bridges to reach the edge of and collectively escape their containers. Comparable cellular systems, such as cytoskeletal walls (Ryan et al., 2012; Simon et al., 2019; Zimmerman et al., 2017) and cellular aggregates (Beaune et al., 2014; Poujade et al., 2007), display protrusion growth that, as in the case of fire ant rafts, facilitates motility and collective migration. While these cellular systems are understood to utilize chemotaxis (Alexandre, 2015), durotaxis (DuChez et al., 2019) or other cue-driven mechanisms (Wen et al., 2015) to initiate migration, it is not entirely clear whether such external stimuli are necessary to drive protrusion growth in condensed active matter and, in particular, fire ant rafts. This raises the question; do fire ants deliberately work to create these protrusions or do these features emerge spontaneously in the absence of targeted signals or external cues? Perhaps more importantly, could such a function be achieved by a condensed system comprised of simple agents interacting through local physical interactions alone? Indeed, Janus particles entrapped by lipid membranes have been shown to generate remarkably similar geometries to these ant rafts due solely to stochastic cooperative motion (Vutukuri et al., 2020). This occurs when a few Janus particles in close proximity simultaneously apply force to the boundary that causes an acute increase in local edge curvature and runaway tether growth. Although the mechanisms driving protrusion growth in ant rafts likely mirrors that of these Janus particle systems, the latter does not constitute condensed active matter, is comprised of two constituent types, and does not display the same dynamic treadmilling mechanisms that replenish ant rafts’ tether growth. Ant rafts, in contrast, are a 2D system comprised entirely of one type of constituent. Furthermore, ant rafts are observed undergoing oscillatory phase changes between highly eccentric periods of growth and relatively inactive states in which the rafts assume more rounded or elliptical shapes (**Fig. 1A-C**). Yet it is not clear what underlying behavioral properties of individual ants drive this global phase change. For these reasons, further investigation of the mechanisms driving ant raft morphology is required. To conduct this investigation, we here develop and employ an ant-inspired, agent-based, numerical model.

## Results

### Modeling ant rafts

Here we overview the discrete numerical model used in this work to contextualize the results presented. Fully detailed derivations and implementation methods are provided in the **Materials and Methods** section. We see in our previous work that the treadmilling of ant rafts is driven by four concurring mechanisms: (1) perpetual contraction of the floating, structural ant network, (2) phase transition of structural ants out of the network into the freely active surface layer, (3) self-propulsion of the freely active surface ants on top of the raft, and (4) phase transition of surface ants into the structural layer at the raft’s edges (**Fig. 1D**). To capture these mechanisms, the model represents ants as discrete agents whose motions are confined to a pseudorandom lattice of water nodes. To represent the biphasic nature of ant rafts, the model consists of a layer of bounded structural agents representing the raft’s structural network, on top of which a layer of unbound surface agents moves (**Fig. 2A-B**). While these two phases behave independently, agents phase transition between them according to a set of ant-inspired rules.

**Figure 2.**
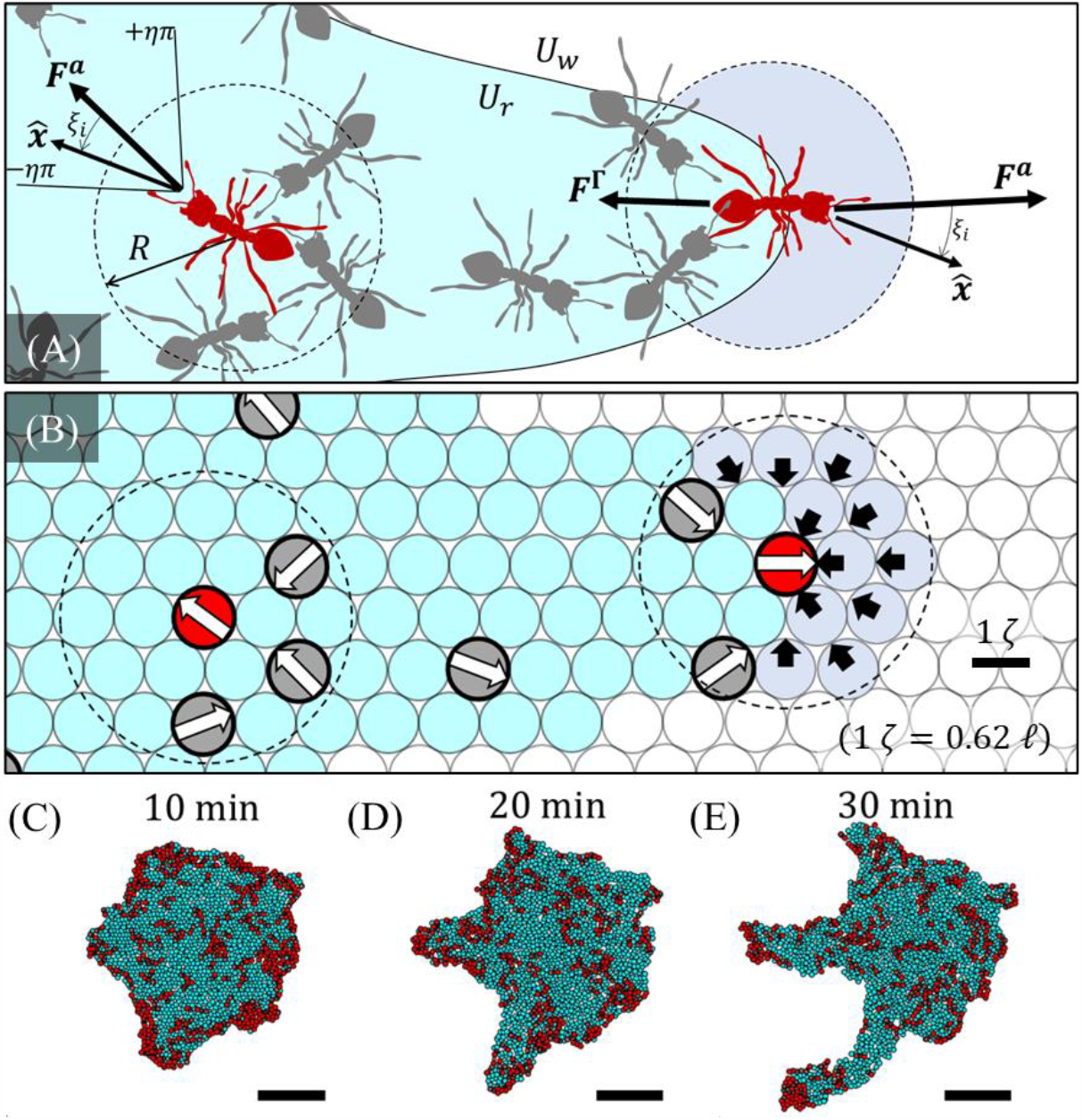
Agent-based Model Schematic. **(A) *F***^***a***^ and ***F***^***Γ***^ are illustrated for two select ants (shaded red): one far from the edge of the raft (left) and one near the edge (right) where the raft is shaded in cyan. ***F***^***a***^ is influenced by the velocities of the *N* neighboring surface ants within detection radius *R*. Likewise, ***F***^***Γ***^ is influenced by the amount and relative position of water within radius *R* (shaded light blue). Note that *R* is not drawn to scale. Trajectory noise due to ant decision-making is introduced via rotation matrix, ***Q***, that redirects surface agents randomly by *ξ*_*i*_ ∈ [**−***ηπ, ηπ*], where the Vicsek noise parameter *η* ∈ [0,1]. **(B)** The numerical implementation of the ABP model defines bound agents (cyan circles) and water nodes (white circles) on which unbound agents (red and grey dipoles) may walk. Motion is updated by identifying the neighboring degree of freedom (DOF) that most closely matches the orientation of intended motion,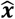. Self-propulsion forces ***f***^***a***^ are depicted as white arrows, while pairwise repulsive forces ***f***^Γ^ from the detected water nodes are depicted as black arrows. Nodes are displayed in a hexagonal, close-packed lattice for illustrative purposes only. **(C-E)** The shape evolution of a simulated raft over a duration of 20 min of simulated time is illustrated and demonstrates protrusion growth. Structural agents are depicted in cyan, while unbound agents are depicted in red. Scale bars represent 10 𝓁.

Based on experimental evidence we find that raft contraction is relatively constant in time (Wagner et al., 2020). In contrast the deposition of surface ants that drives outwards raft expansion varies significantly over hours, with some surface ants clustering near the rod in an inactive state. Therefore, in the scope of this work, our primary aim is to explore the local surface agent behavior that drives phase changes in these systems. Naturally, the model must still replicate the global treadmilling dynamics that are prerequisite to sustained shape change and for which global contraction is an essential mechanism. We found previously that the structural layer contracts uniformly throughout the network and that its density is roughly conserved even over long timeframes (Wagner et al., 2020). To capture this uniform global contraction without introducing mechanisms that would require long-range cooperation, we introduce pair-wise contraction between neighboring structural agents at a constant rate,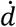. This ensures that the mechanisms driving global contraction could feasibly be achieved by agents working through exclusively local interactions. Setting 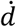 to 1.2 times the globally measured rate contraction rate 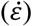 led to good agreement between experiments and simulations (**Fig. 3A-C**) (Wagner et al., 2020). That 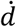 does not equal 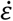 generally indicates that the local rate of contraction between nearest neighbors will be higher than the emergent global rate, which is expected in a network due to non-affine effects (Picu, 2011). To avoid hindering contraction, volume exclusion between structural agents is not enforced. However, unconstrained network contraction would lead to a ceaseless increase in structural ant concentration, which was not observed experimentally (Wagner et al., 2020). To ensure conserved network density structural agents are unbound and converted to active surface agents wherever their local quantity per unit area (*i*.*e*., their density) exceeds a prescribed threshold. To match experiments, this threshold was set to 1 agent per ζ^2^, where ζ^2^ is the area occupied by one experimental, structural ant 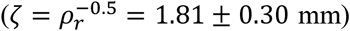. Consequently, a numerical rate of structural unbinding, *δ* ∼ 2 % min^−1^, naturally emerged and mirrored experimental estimates (**Fig. 3F**) (Wagner et al., 2020). Thus, through this pairwise contraction, both global network contraction and flux of ants from the structural network to the surface layer were achieved.

**Figure 3.**
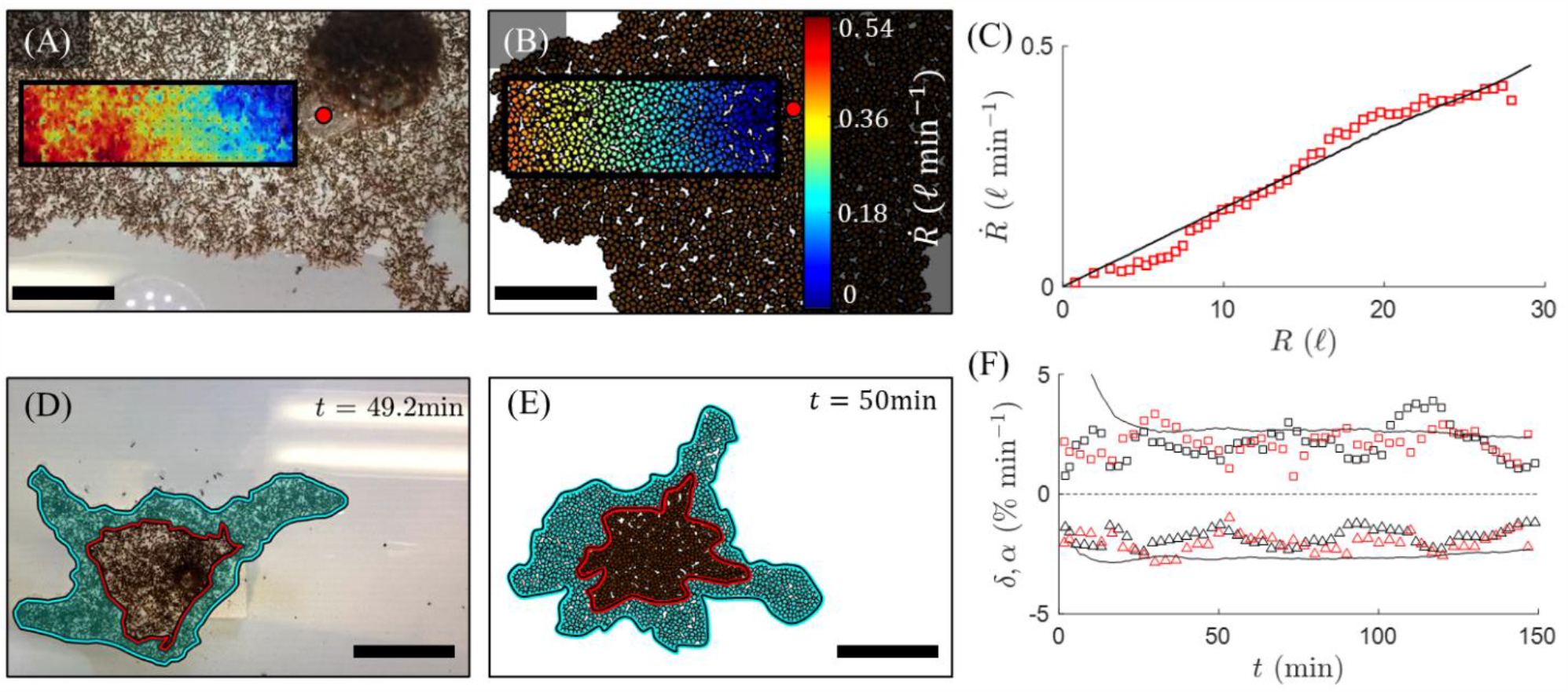
Comparing Treadmilling Dynamics. **(A-B)** The gradient of contractile speed, 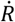, towards the anchor point of the rafts (red dot) is illustrated via heat maps within defined regions of interest (ROIs) for both **(A)** an experimental and **(B)** simulated raft. Scale bars represent 10 𝓁. **(C)** 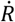 is plotted with respect to distance from the anchor point, *R*, for both the experiment (discrete red squares) and simulation (solid black curve). The slopes of the least-squares regression lines are taken as 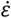. The experimental strain rate 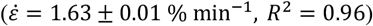 agrees with the numerical value 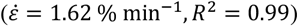. **(D-E)** The growth zones of both **(D)** an experimental and **(E)** simulated raft after roughly 50 min are shaded in cyan. Scale bars represent 15 𝓁. The bound ants that occupied the perimeter of the raft at reference time, *t*_0_ = 0, are outlined in red and were traced through time. **(F)** The time-evolution of the edge binding rate, *α*, and bulk unbinding rate, *δ*, as a percentage per unit raft area are shown for two sets of experiments (squares for *α* and triangles for *δ*; red and black for two different experiments) along with the averaged results of 12 simulations (continuous black curves). Note that the initial drops in both *α* and *δ* for the simulation data occur since the raft was not initiated at steady state, whereas experimental data shown is only taken at pseudo-steady state. **(A**,**C**,**D**,**F)** Experimental results are courtesy of Wagner et al. (2020).

While global contraction and bulk structural unbinding are essential in replenishing the population of freely active ants, it is ultimately the deposition of these free ants into the edge of the structural network that governs global shape evolution. To model surface traffic and edge binding, we began with qualitative observations of freely active surface ants. Namely, that these ants are SPPs whose trajectories under weak confinement are isotropic but correlated below the length scale of one ant body length (1 𝓁) indicating some degree of local alignment (Wagner et al., 2020). Additionally, active ants tend to avoid binding into the raft’s edge, which requires moving into the water, unless pressured by neighboring active ants and adequately surrounded by structural ants upon parking, indicating some aversion to water. We capture the former observation via a local active force ***F***^*a*^, while the latter is addressed by a repulsive boundary force ***F***^Γ^ such that the motion, Δ***x***, of an unbound ant in a discrete timestep, Δ*t*, is well-represented by a reduced form of the generic over-damped Langevin equation typically employed for Active Brownian Particles (ABP) (Shaebani et al., 2020):

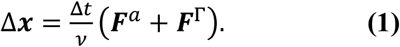

Here *v* is an overdamping frictional coefficient constraining the speed of agents and the environmentally driven diffusion captured by the ABP model is embedded in ***F***^***a***^ through local interactions (see **Materials and Methods** for details). The timestep is defined as Δ*t* = ⟨*d*⟩/*v*_0_, where ⟨*d*⟩ is the average distance an agent will travel on the lattice in one iteration and *v*_0_ is the experimentally measured surface ant speed.

To compute ***F***^***a***^ we enforce that surface agents are driven by their own active force, 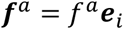, with constant magnitude *f*^*a*^, in the direction of unit vector ***e***_***i***_ (De Magistris and Marenduzzo, 2015; Shaebani et al., 2020). Since we previously observed correlated motion between ants separated by a distance of less than 1 𝓁 (indicative of mutual influence within some radius *R* ∼ 1 𝓁) (Wagner et al., 2020), we take ***F***^***a***^ as the combined effect of each *i*^*th*^ agent’s self-propulsion and that of its *N* neighbors (within radius *R*) such that ***F***^*a*^ = *f*^*a*^***Q*** ⋅ [*N*^**−**1^ ∑_*j*∈[1,*N*]_ ***e***_***j***_]. Rotation matrix ***Q*** induces a random change in agents’ directions in the range [*πη, πη*], thus capturing decision-based noise in ants’ trajectories. Here *η* ∈ [0,1] is the noise parameter introduced by Vicsek et al. (1995) (**Fig. 2A**).

Recalling surface ants’ observed aversion to water, we computed ***F***^Γ^ as the spatial gradient of an effective potential field, **−**∇_*r*_*U*, that the agents enter when they detect water (again within detection radius *R*). ***F***^Γ^ is not a physical force, but rather an embodiment of ants’ motivation to stay on dry substrates and is akin to the “social forces” employed by Helbing and Molnár (1995). First, we designate potentials *U*_*w*_ and *U*_*r*_ that represent unbound agents’ affinities towards water nodes and structural agents, respectively (*U*_*r*_ > *U*_*w*_). The result is a pair-wise potential force, 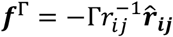, between each *i*^*th*^ surface agent and its *j*^*th*^ water neighbor, where Γ = *U*_*w*_ **−** *U*_*r*_ < 0 and *r*_*ij*_ is the separation vector between their positions. Summing these pair-wise forces over all *N* detected water neighbors gives 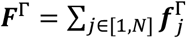. In effect, ***F***^Γ^ repels unbound surface agents from the water proportionately to the amount of water detected (or “felt’’ since *R* ∼ 1 𝓁). This occurs approximately in the direction normal to the raft’s boundary.

When agents are far from the raft edge (***F***^Γ^ = 0), **Equation 1** degenerates to the form of Vicsek model employed by Hugues Chaté et al. (2008), predicting that free ants walk at a speed *v*_0_ ≈ *f*^*a*^/*v* with an average persistence length *l*_*p*_, which depends on *η* and *R*, taken as 0.2 and 0.9 𝓁, respectively (Vicsek et al., 1995). However, at the edge of the raft, according to **Equation 1**, the condition for agent parking is dictated by the net force along its direction of motion, 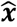, as 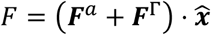. Our algorithm enforced structural edge binding only if *F* > 0. With this rule implemented, a global expansion (or edge binding) rate *α* ∼ 2 % min^−1^ naturally emerged and automatically matched the experimentally measured values once pseudo-steady state treadmilling occurred (*α* ≈ *δ*) (**Fig. 3**.**D-F**). We found that edge binding depends on a single length scale *L* = Γ/*f*^*a*^, interpreted as the travel distance over which the work of one ant (given by *f*^*a*^*L*) is comparable to the edge energy barrier (given by Γ). Taking Γ as a constant (*i*.*e*., assuming ants’ aversion to water does not change), reducing *L* implies that the surface agents self-propel with a greater magnitude of force *f*^*a*^ and that they do the same amount of work in less distance (*i*.*e*., smaller *L*). Normalizing the body length of an ant by *L*, we defined a dimensionless activity parameter given by:

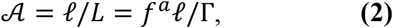

that measures the relative energy of an unbound surface agent as compared to the edge repulsion potential.

### Protrusions emerge spontaneously

To mirror experiments, we ran simulations with 2,250 agents for up to 4.5 hours of simulation time, letting the rafts reach quasi-steady state treadmilling (wherein *α* ≈ *δ*). To roughly mimic the dense, spheroidal shapes of the ant aggregations at initial time and to assure controlled initial conditions, the simulated rafts were originated as circles with a surface packing fraction of *ϕ* = 1 surface agents per structural agent (**Fig. 5**). To provide a still reference frame and mimic the anchored boundary condition of experimental rafts, a permanent structural agent was located at the center of the domain and fixed in place. With both pairwise contraction rates 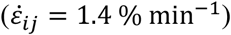 and the Vicsek model parameters (*R* = 0.9 𝓁 and *η* = 0.2) independently calibrated to match experimental treadmilling and free ant trajectories, respectively, ant activity (𝒜) remained the only free parameter driving surface agent behavior and global shape evolution. When 𝒜 was on the order of 1.25 to 1.47 the model predicted the unstable growth of protrusions. To determine if the predicted protrusions had the same characteristic length scale and dynamics as experimental protrusions, we measured their average widths, *W*, and growth rates, 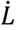, over time. The widths of simulated protrusions ranged from roughly 2 to 9 𝓁, with a mean value of 5.95 ± 0.05 𝓁 that agrees with the experimental value of 5.85 ± 0.06 𝓁 (**Fig. 4A**) (Wagner et al., 2020). Similarly, the tip-growth rates of the model-predicted protrusions ranged from roughly −1 to 3 𝓁 min^−1^, with a mean value of 0.46 ± 0.02 𝓁 min^**−**1^ (**Fig. 4B**). While not exactly matching the experimental mean of 0.74 ± 0.05 𝓁 min^**−**1^ (Wagner et al., 2020), these growth rates are on the same order and are adjustable through 𝒜. Our model also allowed us to easily quantify the distinct behaviors of unbound agents, enabling us to confirm the factors leading to spontaneous protrusion initiation and runaway growth.

**Figure 4.**
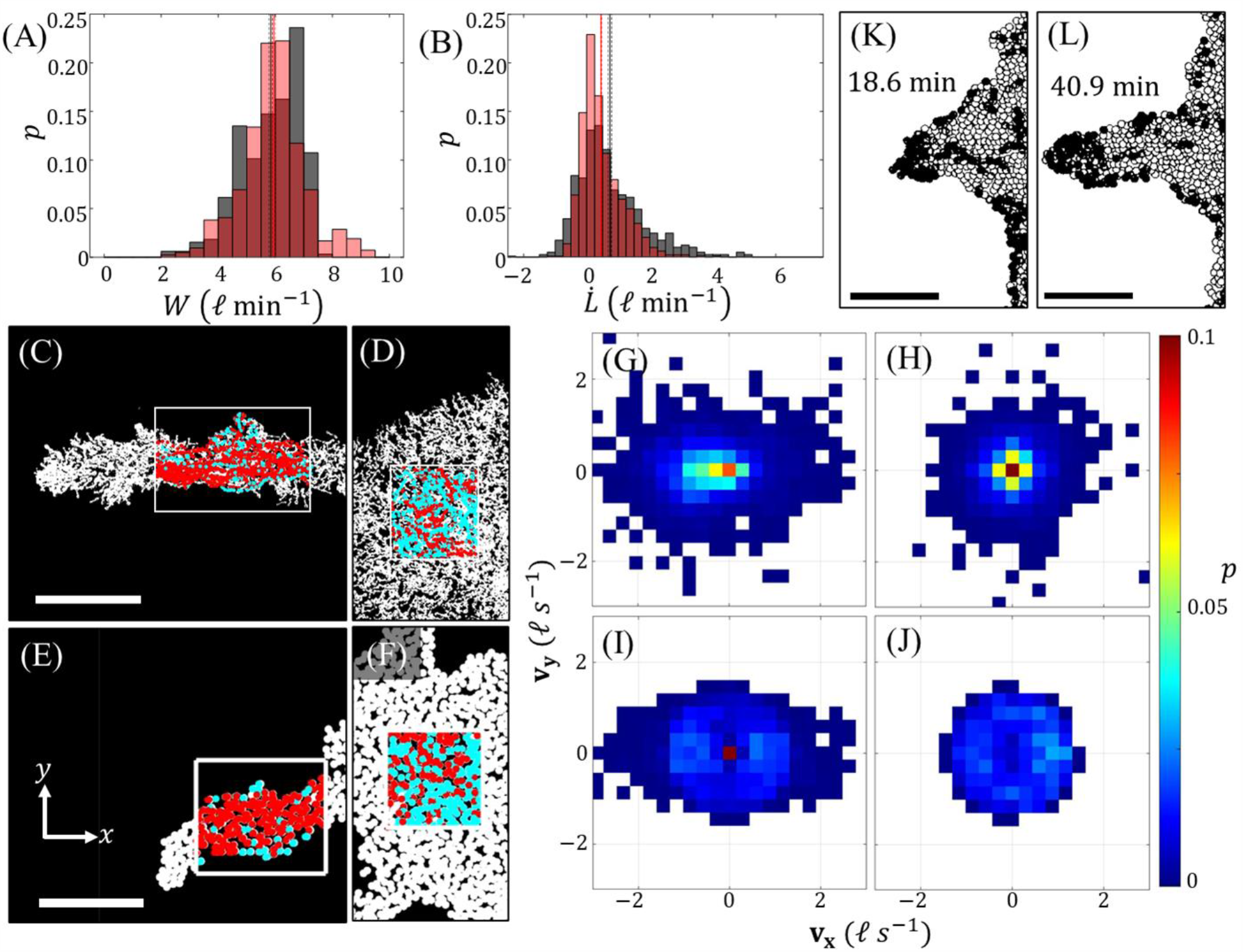
Comparing Protrusion Dynamics. **(A-B)** Probability distribution histograms are shown for (**A**) the average protrusion widths, *W*, and (**B**) growths rates, 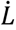 of more than 400 experimental observations (grey) and numerical observations (light red) each. Here, *R* = 0.9 𝓁, *η* = 0.2 and 𝒜 ∈ [1.25,1.47]. **(C-D)** The normalized order parameter *φ* for (**C**) a section of experimental raft on the protrusion is higher than that of (**D**) a section on the bulk of the raft. Note that, *φ* → 0 indicates disordered motion, whereas *φ* = 1 indicates completely synchronous motion. **(E-F)** The same comparison is made for (**E**) a section of protrusion to that of (**F**) a section on the bulk for a simulated raft. For **(C-F)** leftwards moving unbound ants are illustrated in red, while rightwards moving ants are illustrated in cyan to clearly illustrate the anisotropic, directional flow that frequently occurs on protrusions. (**G-J**) The 2D probability velocity distributions corresponding to (**C-F**), respectively, are shown. A simulated protrusion at the start **(K)** and end **(L)** of a roughly 21 min duration exhibits how directional motion on protrusions culminates in clustering of surface agents (black circles) at the tip and rapid, anisotropic growth. **(A-B**,**C-D**,**G-H)** Experimental results are courtesy of Wagner et al. (2020). Scale bars in (**C**,**E**,**K**,**L**) represent 10 𝓁.

**Figure 5.**
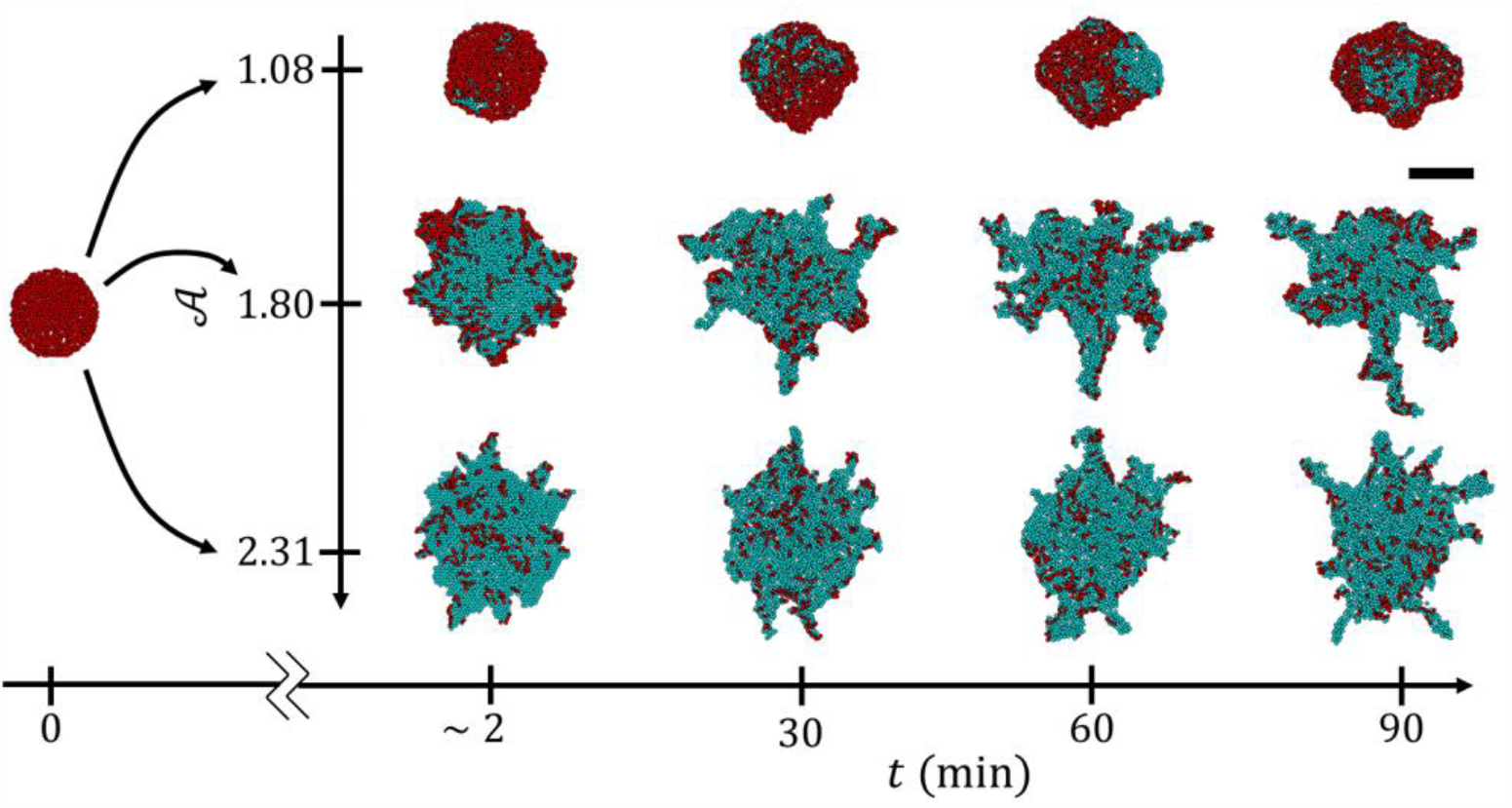
Dynamic Effects of Activity Level. Snapshots of modelled rafts at different simulation times (*t*) and activity levels (𝒜) are depicted to illustrate the effect of 𝒜 on raft development. The raft on the far left depicts the initial conditions which were the same for each simulation (a circular raft with *ϕ* = 1). On the right each row depicts a single raft as it evolves in time (moving from left to right along the horizontal axis). For all simulations shown, *η* = 0.2 and *R* = 0.9 𝓁. Structural agents are depicted in cyan, while unbound agents are depicted in red. The scale bar in the top right is universal to all snapshots and represents 14 𝓁.

We confirmed that protrusion initiation is driven by stochastic nucleation of ant clusters, which occurred frequently near the rafts’ convex edges due to wall-accumulation effects (Dong et al., 1994; Elgeti and Gompper, 2013; Fily et al., 2014; Méhes and Vicsek, 2014; Reichhardt et al., 2018; Shaebani et al., 2020). These clusters often caused concentrated edge-deposition of unbound ants resulting in local regions of high edge curvature that served as proto-protrusions. Following this, the model predicted directional flow of unbound ants along protrusions’ lengths consistent with what was observed previously in experiments (**Fig. 4C-F**,) (Wagner et al., 2020). We quantified this by comparing the normalized velocity order parameter of surface ants and agents (members) on protrusions, defined by *φ* = ⟨***v***(*t*)⟩_*N*_/⟨|***v***(*t*)|⟩_*N*_ where ***v*** is their velocity of a particle and ⟨⟩_*N*_ denotes taking the ensemble average over all *N* members (Vicsek and Zafeiris, 2012). This parameter is zero when motion is completely isotropic but approaches 1 when movement is perfectly unidirectional. We found that in both experiments and simulations, *φ* was consistently higher for surface members on protrusions than on the bulk of the raft. For the experimental raft depicted in **Fig. 4C-D**, *φ* = 0.65 ± 0.02 on the protrusion versus *φ* = 0.57 ± 0.02 on the bulk (Wagner et al., 2020). Similarly, for the simulated raft depicted in **Fig. 4E-F**, *φ* = 0.64 ± 0.12 on the protrusion versus *φ* = 0.33 ± 0.06 on the bulk. In both cases, comparably sized domains were used to compute *φ* and the relatively larger values of *φ* on protrusions confirms that the confinement of protrusions induces higher directional motion than that observed on the bulk. However, *φ* does not indicate the orientation or sense of said directional motion. To examine in which orientation surface members preferentially traveled, we also investigated their velocity distributions on and off protrusions, from both experiments (**Fig. 4**.**G-H**) (Wagner et al., 2020) and simulations (**Fig. 4**.**I-J**). The elongation of velocity distributions along the length of protrusions confirms that traffic moves primarily along these growths’ longitudinal axes. To visually illustrate the sense of this motion, all surface members moving left (from base-to-tip of the protrusions) are overlaid with red in **Fig. 4C-F**, whereas members moving right are overlaid with cyan. The relatively higher presence of red in **Fig. 4C** and **4E** qualitatively exhibits that at the time of investigation, most of the surface members were moving from base-to-tip of the protrusions. When directional traffic occurred towards the tips of protrusions, it induced jamming of unbound agents at their ends (**Fig. 4**.**K-L**) and high magnitudes of ***F***^***a***^, similar to the locally high pressures exerted by confined Active Brownian Particles on regions of highly convex wall curvature (Fily et al., 2014; Nikola et al., 2016). This locally high active force accentuated edge binding and tip growth. Ultimately, runaway protrusions result from a positive feedback loop wherein cluster formations initiate protrusions that in turn promote directional traffic, spurring further tip clustering and growth. Indefinite growth of protrusions is checked by both the perpetual raft contraction and finite population of surface agents, such that within an appropriately large domain the protrusions do not reach the simulation’s boundaries.

### Activity level modulates shape

Having confirmed that local-level agent interactions can lead to spontaneous instabilities, we then sought to understand the local behavioral changes that could lead to long-term variation in experimentally observed raft shapes by exploring the effects of 𝒜 over the range of 0.81 to 3.23 (see **Movies S2-S9** for samples in this range). Results are summarized in **Fig 5** and **Fig. 6A** where we visually present the effects of activity on the raft configurations during and after 1.5 hours of simulated time, respectively. From **Fig. 5**, we see that no protrusions emerged when 𝒜 = 1.08, whereas protrusions emerged within the first 30 min and 2 min when 𝒜 = 1.80 and 2.31, respectively. Similarly, while there are numerous protrusions stemming from the rafts in **Fig. 6A** when 𝒜 ≥ 1.47, there were no distinct growths on the rafts when 𝒜 ≤ 1.16. Generally, these results indicate that higher 𝒜 promotes higher parking rates that induced more frequent protrusion growth and generally higher surface excess.

**Figure 6.**
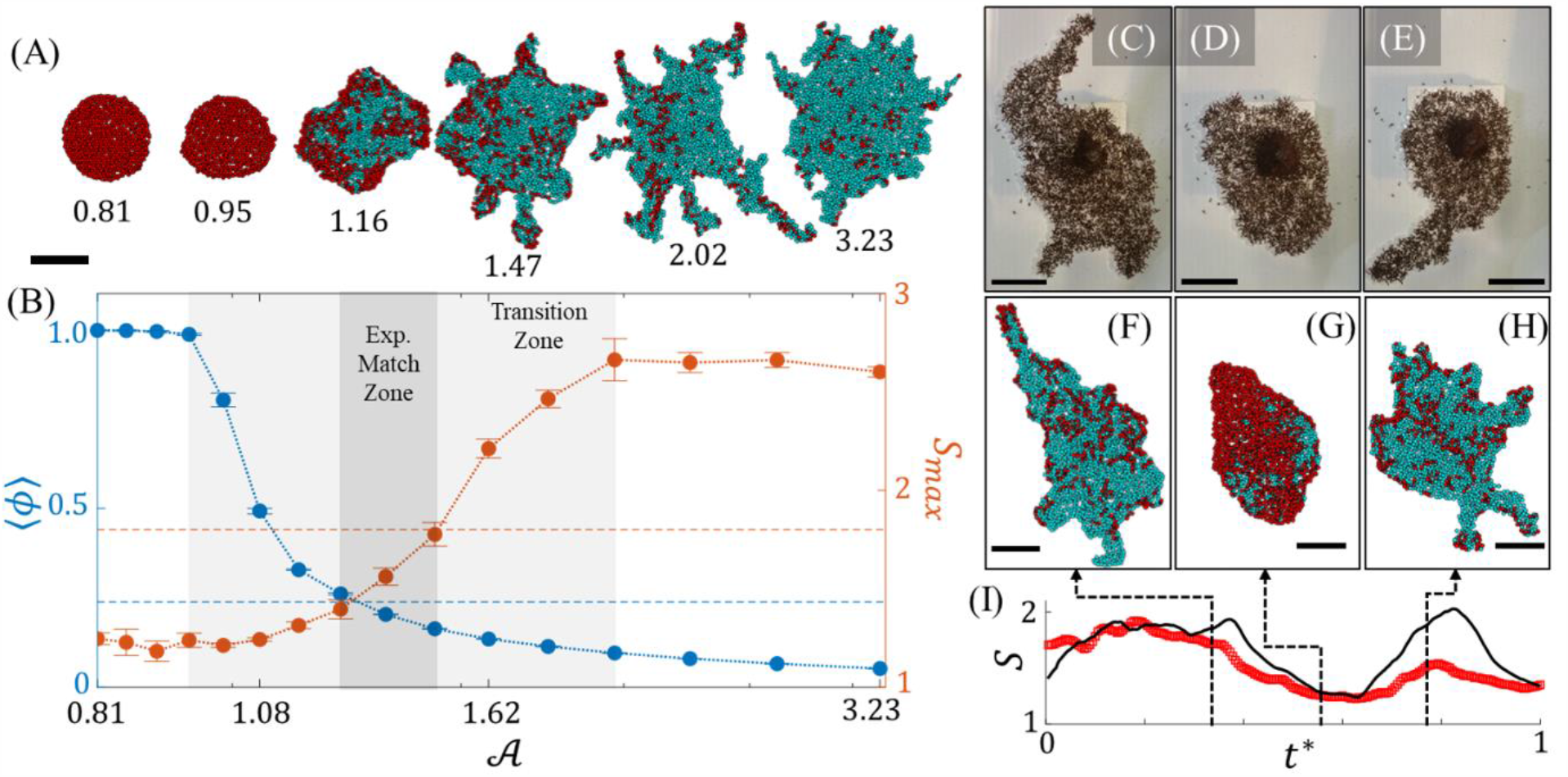
Ant Activity Phases. **(A)** Snapshots of simulated rafts after 1.5 hours of simulation time are shown for which *η* = 0.2, *R* = 0.9 𝓁 and 𝒜 ∈ [0.81,3.23]. The value of 𝒜 yielding each morphology is denoted beneath each snapshot. **(B)** Mean unbound agent packing fraction, ⟨*ϕ*⟩, (blue) and maximum surface excess, *S*_*max*_, (red) are plotted with respect to 𝒜 averaged over 5 simulations at each value of 𝒜, with error bars presenting standard error. Horizontal dotted lines in **(B)** represent the manually experimentally measured values of ⟨*ϕ*⟩ = 0.24 and *S*_*max*_ ≈ 1.8. The bounds of the parameter space that matches our experiments are marked where these respective lines intersect the numerical data (see “Exp. Match Zone” between 𝒜 = 1.25 and 1.47). There exists a zone between roughly 𝒜 = 1.0 and 2.0 of continuous phase transition between rafts with minimal-to-no growth whatsoever (*ϕ* ≈ 1 and *S* ∼ 1.2) at low activity levels and frequent protrusion growth (low *ϕ* and *S* > 2) at high activity levels. **(C-H)** Three chronological snapshots of an experimental ant raft (**C-E**) exhibiting different phases of protrusion growth are compared to three chronological snapshots of a simulated raft (**F-H**) when 𝒜 was modulated between 1.1 and 1.6. **(I)** The time evolution of surface excess from the experiment (red squares) and simulation (black curve) are displayed. Time, *t*^∗^, is normalized by the experiment duration for a more direct comparison. Structural agents are depicted in cyan, while unbound agents are depicted in red. All scale bars represent 10 𝓁.

To quantitatively characterize global shape, we introduce a dimensionless parameter called surface excess defined by 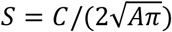, where *C* and *A* are a raft’s perimeter length and area, respectively. *S* = 1 for a circle and increases with a shape’s eccentricity, thus higher *S* indicates the presence of more numerous or more elongated protrusions. Maximum surface excess and mean surface packing fractions of model results are presented in **Fig. 6B**. Maximum (as opposed to mean) surface excess is presented to transparently indicate the peak degree of eccentricity achieved by the raft and exclude the inherently low surface excess of the initially circular rafts. Moving from left to right in **Fig. 6B** there is a continuous phase transition in the activity range of 𝒜 = 0.95 to 2.02 indicated by the smooth curve of surface excess from low (*S* ≈ 1.2) to high (*S* ≈ 2.8). Likewise, there is a smooth transition of unbound packing fraction from high (*ϕ* ≈ 1 implying almost no edge binding, whatsoever) to low (*ϕ* ≈ 0.06, indicating relatively high edge binding rates) as 𝒜 increases. These phases are analogous to those observed in **Fig. 1A** and **Fig. 1B**, respectively.

To demonstrate how 𝒜 may alter the phases of protrusion growth and non-growth, we modulated 𝒜 within a given model experiment to a value above (𝒜 = 1.6) and below (𝒜 = 1.1) the phase transition threshold (see **Fig. 6**.**C-I** and **Movie S12**). Indeed, this effectively toggled the raft between exploratory phases of high surface excess (**Fig. 6F** and **6H**) and low surface excess wherein no protrusions were present (**Fig. 6G**), comparable to what was observed in experiments when unbound surface ants ceased activity. Worth noting is that the second phase of experimental protrusion growth (**Fig. 6E**) did not reach the same magnitude of surface excess as the initial phase (**Fig. 6C**), suggesting that either the activity level did not fully recover to its original state or not enough time was spent in this more active state to resume *S* ≈ 2. Consequently, surface excess of the second simulated phase of protrusion growth (**Fig. 6H**), exceeds that of the experimental surface excess. It is reasonable to assume that 𝒜 will evolve roughly continuously for a real ant raft of thousands of individuals, however 𝒜 in the model was modulated via a binary step function, thus likely contributing to the abrupt resumption of high *S*. Regardless, activity level’s effect on raft shape is made clear.

Besides dictating the global presence of protrusions, 𝒜 also impacts the local shape of these tethers. Specifically, higher 𝒜 diminishes the characteristic widths of growths. Mlot et al. (2011) estimated the capillary length of ant rafts on the order of 10 𝓁. However, here we see that raft edge curvature is dependent on the length scale, *L* = 𝓁/𝒜, and therefore mediated by activity. When 𝒜 is low, *L* increases, and we see smoother raft geometries (higher capillary length) with lower surface excess. In contrast, when 𝒜 is high, *L* diminishes permitting the emergence of more, but narrower, protrusions. In essence, higher unbound agent activity reduces effective surface tension of the overall rafts, warranting a comparison of 𝒜 to temperature (Loi et al., 2008) in non-active materials whose surface tensions generally diminish as temperature increases (Cini et al., 1972). Worth noting is that when 𝒜 is sufficiently high, the rate and azimuthal homogeneity of edge binding appears significantly high such that expansion appears to approach an isotropic state. This is reflected by the reduction in the number of and size of protrusions displayed by the raft in **Fig. 6A** when 𝒜 = 3.23. It is likely that as 𝒜 increases, the propulsion force of a single agent ***f***^***a***^ eventually becomes sufficient to cause edge binding anywhere along the raft’s edge, reducing the relative significance of local raft geometry and cooperative force ***F***^***a***^. Significantly, this suggests that there is, in fact, an optimal surface activity level for inducing an exploratory phase in systems that obey this model, which occurs when 1.16 < 𝒜 < 3.23.

## Discussion

Our model indicates that fire ant rafts may exhibit spontaneous protrusion growth in the absence of external cues, long-range interactions, or centralized control and that the global response of these condensed active systems depends on the underlying behavior of individual constituents. Specifically, we find that the global shape of these rafts is highly dependent on the activity parameter – a dimensionless parameter that characterizes the competition between an ant’s self-propulsion and its aversion to water. Tennenbaum and Fernandez-Nieves (2020) demonstrated that temporal activity cycles in fire ants on the order of hours impact the rheological properties of 3D aggregations. We expect that this same oscillation of ant activity level is responsible for the morphological variation in ant rafts, however further experimental work is required to quantitatively confirm that there is, in fact, an increase in surface activity during periods of high outwards expansion.

While this work provides a potential explanation for protrusion emergence through exclusively local interactions, we do not claim to have ruled out biological stimuli (*e*.*g*., morphogens or pheromones). Instead, we recognize that our ant-inspired model may have identified a redundant set of local mechanisms through which fire ants could feasibly achieve this emergent phenomenon spontaneously. Our results may suggest that there exists redundant pathways or physics-driven mechanisms that induce emergent protrusion growth in other biphasic active matter systems as well, perhaps including cytoskeletons and cellular aggregates (Bugyi and Carlier, 2010; Méhes and Vicsek, 2014; Neuhaus et al., 1983), even in the absence of biological stimuli. Therefore, although this numerical implementation was inspired by fire ant rafts, we expect that in future work it may be adapted in or inspiration for the *in silico* investigation of other biological systems driven by transport and binding reactions. These results may also inform the predictive design of engineered systems such as active gels or swarm robotics. For example, by altering the self-propulsion activity of individuals in cohered robotic swarms, researchers could potentially induce a phase of collective expansion which can be utilized for group exploration.

## Materials and Methods

### Model Derivation and Implementation

Here we detail the derivation and numerical implementation of the model used to simulate both structural and surface ants as discrete agents. In the remainder of this section, we use the index, *i*, to denote the agent of interest, and *j* to denote the influencing neighbors of *i*.

### Domain Description

Our numerical framework is carried out using MATLAB R20109b. It is a 2D planar, discrete model comprised of distinct nodes defined by some unique index number, *i* ∈ [1: ∞), and unique Cartesian coordinates, ***X***_*i*_ = [*x*_*i*_, *y*_*i*_]. We locate the nodes inside of a square domain whose center is at position [0,0]. The nodes are initially positioned in a close-packed hexagonal lattice with unit length spacing between nearest neighbors. Each node position is then offset by some random amount in the range [**−**1/6: 1/6]. At initial time, *t*_0_ = 0, each node is classified as either a structural agent (shown as cyan circles in **Fig. S1**), or water node (shown as black dots in **Fig. S1**). Structural agents represent structural ants, whereas water nodes represent vacant locations into which ants may eventually park. To mimic initial experimental conditions, the initial raft structure is a circle with center [0,0] and some prescribed radius (**Fig. 5**). Every node within this circular boundary is initially defined as a structural agent. This ensures that any protrusions predicted by the model emerge spontaneously as opposed to through artificial asymmetries. Surface agents are introduced to represent surface ants (shown as red circles in **Fig. S1**). The initial surface packing fraction is set to 1 surface ant per raft ant. To simulate the movement of surface ants on top of the raft, we require that these surface ants only occupy sites already designated as raft nodes. Additionally, to simplify the model, we enforce volume exclusion between surface agents so no two surface agents to occupy the same raft site.

### Length Scales

Two length scales are referenced in this work. The first is that of the mean ant body length, taken as 1 𝓁 = 2.93 mm. Results are generally presented in this length scale for ease of comprehension and comparison to experimental results. However, the numerical model is normalized by a second length scale defined as 1 ζ = 1.81 mm. 1 ζ^2^ is defined as the area a single ant in the structural raft network occupies (*i*.*e*., 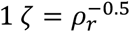, where *ρ_r_* = 0.304 ants mm^**−**2^ is the density of structural ants). This normalization permits that the density of structural agents may be maintained at 1 node per unit area (ζ^2^), and the nominal separation between nodes will be on the order of 1 unit length (ζ).

### Time Scale

We normalize the timescale by taking the average distance a surface agent will travel in one iteration, ⟨*d*⟩, divided by the average experimentally measured surface ant speed, *v*_0_ (*i*.*e*., Δ*t* = ⟨*d*⟩/*v*_0_).

**Figure S1.**
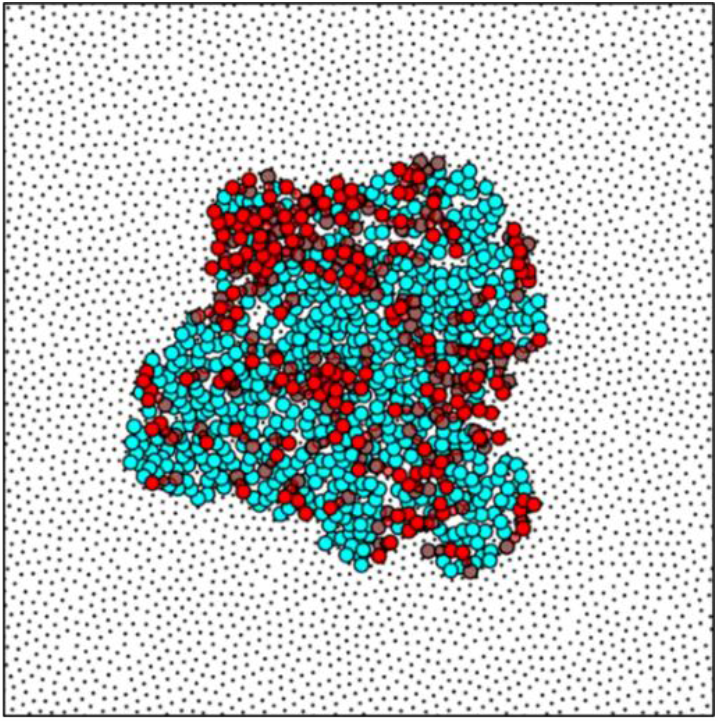
A snapshot of the discrete numerical domain is shown with water nodes plotted as black dots, raft nodes plotted as cyan circles and surface agents plotted as red circles.

### Simulating the Structural Network

Given the experimentally measured contraction rate 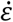, we apply a pair-wise strain rate, 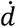, between connected neighbors within the raft. Through 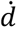, we can calculate the updated pairwise separation vector, ***d***_*ij*_, between all raft nodes and their raft neighbors residing within some prescribed radius, *R*_*r*_, about the node of interest. We conservatively set to *R*_*r*_ to 1.5 𝓁 to capture the bridging of raft voids that often occurs between structural raft ants. *R*_*r*_ = 1.5 𝓁 corresponds to roughly 4.5 mm or half the body length of some of the largest fire ants. To implement raft contraction, we enforce that the separation distance between all raft neighbors at time *t* decays according to:

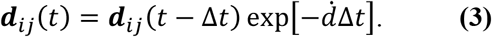

Taking ***d***_*ij*_ as the targeted equilibrium separation at time, *t*, we then define the required displacement as:

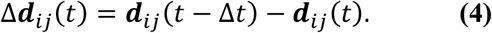

***X***_*i*_ for each raft node is then updated according to:

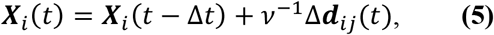

where *v* ∈ [1, ∞) is simply a pseudo-viscosity or over-damping scalar used for computational stability. Given the new ***X***_*i*_, we then re-calculate ***d***_*ij*_, take it as the updated ***d***_*ij*_(*t* **−** Δ*t*) (in **Equation 3**), and iterate **Equation 4** and **Equation 5** until the residuals dip below some prescribed threshold. Here we define the residuals and their thresholds as max[Δ*d*_*ij*_(*t*)] ≤ 5 × 10^**−**5^ ζ and mean[Δ*d*_*ij*_(*t*)] ≤ 1 × 10^**−**5^ ζ.

### Close-packing Water Nodes

Given that the raft will shrink in time, we need to ensure that water nodes will remain closely packed to the edges of the raft and binding events on the edge remain possible. To do this, we apply a radial linear velocity gradient to all water nodes that moves them towards the center of the domain, [0,0], at the rate of 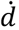, according to 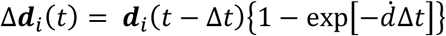, where ***d***_*i*_ is the separation vector of each water node with respect to [0,0]. To evenly space water nodes from each other, as well as the raft edges, we introduce Gaussian, pair-wise repulsive potentials, 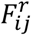, between water nodes and their nearest raft or water neighbors of the form:

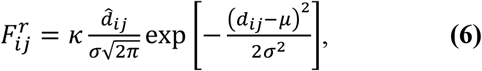

where σ is the standard deviation of the curve, μ = 0 ζ is the mean, and *κ* is a scaling factor in units of pseudo-force. We use these values to step the displacement of water nodes according to:

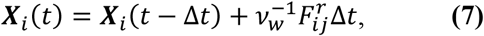

where *v*_*w*_ ∈ [1, ∞) is simply another pseudo-viscosity for computational stability. We found that σ = 0.5 ζ and *k*/*v*_*w*_ = 0.02 provided a stable computational domain in which water nodes remained close-packed in an evenly distributed point field, thus offering ample water degrees of freedom, for surface agents on the edge of the raft to park into. Note that the repulsive interactions between water nodes and their raft neighbors were one-way such that the raft nodes displaced water nodes, but water nodes could not move their raft neighbors. This was done because the close packing of water nodes is a numerical method implemented to homogenize the domain rather than any physical phenomena and should not influence the position of raft nodes in our model.

### Unbinding to Maintain Structural Density

Recall that we normalize the domain’s unit length by ζ such that the domain’s nominal density, *ρ_d_*, is approximately 1 node ζ^**−**2^. Since we observed that structural network density remains roughly constant, we enforce unbinding events in simulations when the domain density exceeds 1 node ζ^**−**2^. To ensure that nodes are removed from the densest locations with precision, we subdivide the domain into a square grid whose unit cell lengths, *L*_*g*_, are ≥ 1ζ. The number of permissible nodes, *N*_*p*_, within each grid cell becomes 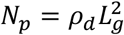. For our purposes, we found that *L*_g_ = 2ζ and *N^p^* = 4 nodes provided sufficient regional discretion in order to maintain a homogeneous domain (see black dots in **Fig. S1**).

At every time step, we conduct a count of the number of nodes occupying each grid site and if the count exceeds *N*_*p*_, we initiate a node deletion event. To introduce further specificity in which nodes to remove we calculate the pair-wise distance, *d*_*ij*_, between each node in the pertinent grid space. If both nodes *i* and *j* belonging to the smallest value of *d*_*ij*_ occupy the grid space, one of the two is randomly selected for deletion. In the case that the deleted node is a water node, we simply delete it. However, if the deleted node is occupied by a structural agent, we convert it to a surface agent positioned at the coordinates of the nearest empty structural node. By counting the number of unbinding events at each time step and normalizing by the area of the raft, *A* or *N*_*r*_/*ρ_r_*, we can calculate the unbinding rate according to *δ* = *N*_*r*_/(*ρ_r_*Δ*t*), where *N*_*r*_ total number of structural raft agents and *ρ_r_* = 1 node ζ^**−**2^.

### Simulating the Surface Ants

Surface agents are only permitted to occupy the discrete positions already occupied by structural raft agents. To enforce volume exclusion, no more than one surface agent is permitted to occupy a given position at the same time. Thus, while water and structural agents exist within the continuous 2D domain, surface agents may be thought of as existing on top of the lattice comprised of these other nodes.

To model the active surface ants, we assign every node in the domain 18 degrees of freedom (DOF), which – in an equidistant hexagonal lattice of nominal unit separation – corresponds to the number of nearest neighbors spanning 2 ζ. We opted to provide each node with 18 DOF based on three considerations: (1) the experimental observation that surface ants frequently walk over one another, effectively passing 2 ζ in one unit time; (2) the experimental observation that surface ants may walk over voids in the raft of comparable dimensions to their own body length, effectively passing greater than 1 ζ in one unit time; and (3) the realization that modeling surface agents with 18 rotational DOF is roughly a threefold improvement in approximating the continuous space real ants occupy, over the 6 DOF offered by looking at only one layer of immediate node neighbors spanning 1 ζ. The 18 DOF for each node are assigned in rank-order by distance to neighboring nodes. To be a DOF, the neighboring node must reside within the distance, *R*_*DoF*_ ∈ (0,2.5] ζ, of the node of interest. The upper bound of this range was set to 1.25 × the distance needed to reach 18 nearest neighbors in a close-packed hexagonal lattice, to account for noise.

We define the velocities of each node at time *t* as ***v***_*i*_(*t*) = [***X***_*i*_(*t*) **− *X***_*i*_(*t* **−** Δ*t*)]/Δ*t*. Note that the magnitude of the particle velocity, |***v***_*i*_|, is approximately limited by the range |***v***_*i*_| ∈ (0,2.5 ζ/Δ*t*] due to the distance limits over which DOF are considered. Therefore, we assign the system a velocity modulus, *v*_0_, which is simply the experimentally measured average ant speed. Assuming all 18 DOF are equally probable movement locations in a hexagonal lattice, then the average magnitude of movement between adjacent neighbors is, ⟨*d*⟩ = 1.67 ζ, and the normalized timescale (recall, Δ*t* = ⟨*d*⟩/*v*_0_) comes out to roughly Δ*t* = 1.7 s per iteration. Note that the velocities of surface agents in our model are dependent on those of the previous time step, so we must kick-start the system through assignment of random initial velocities.

### Stepping the Movement

To kick-start the movement of surface ants at the start of our simulation or when a structural agent transitions to a surface agent, we assign each surface agent a random, instantaneous velocity orientation, *θ*_*i*_ ∈ [0,2*π*] radians, as measured from the positive horizontal axis. In this quasi-lattice framework, each surface agent has at most 18 DOF that can be classified as unoccupied nodes on the raft, occupied nodes on the raft, or water nodes. To determine which DOF each surface agent preferentially moves to at each time step, we predict the preferred angle of movement, *θ*_*i*_, using the over-damped Langevin equations commonly used for Active Brownian Particles (Shaebani et al., 2020), detailed in the following section, **Equations of Motion**.

With the preferred angle of motion predicted, we identify the neighboring DOF whose orientation with respect to the surface agent’s position is closest to this preferred direction. We calculate the relative angle of each DOF, ϑ_*ij*_, as measured with respect to the positive horizontal axis according to:

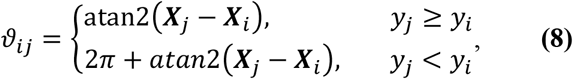

where ***X***_*j*_ is the position of each DOF and ***X***_*i*_ is position of the surface agent. We then calculate the absolute difference between *θ*_*i*_ and ϑ_*ij*_:

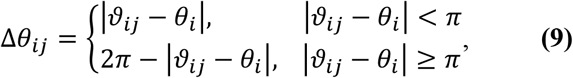

and take the minimum value to indicate which neighboring raft node or water node the surface agent would preferentially move to. The DOF are then rank ordered from smallest to largest Δ*θ*_*ij*_ into a pool of potential movement nodes and sequentially checked for eligibility of movement. Note that to be eligible for movement, neighboring raft and water nodes must exist within the confines of some turning limit (*θ*_*l*_ ∼ ±0.5*π* radians). This turning limit was included due to the observation that it takes the ants greater than Δ*t* to turn more than approximately *π*/2 radians.

With the rank ordered DOF, we then sequentially check for parking eligibility according to the following section. The algorithmic chronology used for determining movement of surface agent *i* is depicted in **Fig. S2**.

**Figure S2.**
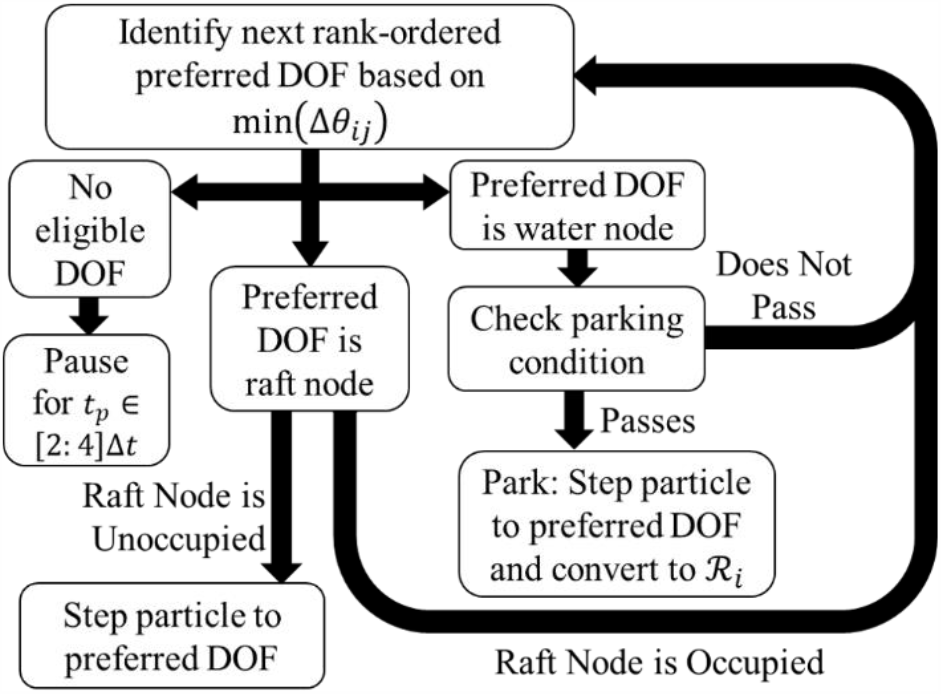
Algorithmic Chronology: A flow chart details the algorithm by which movement is determined for each surface agent, in each time step. Here, ℛ_*i*_ denotes a structural agent.

### Equations of Motion

We determine the preferred direction of motion using the over damped Langevin equations of motion (Shaebani et al., 2020) that give the velocity, ***v***_*i*_, and rate change of particle direction, 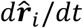, of a particle as:

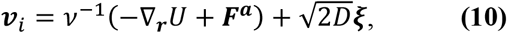

and:

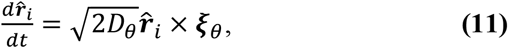

respectively. Here, *v* is the overdamping or frictional coefficient; *D* is the translational diffusion coefficient; *U* is the local potential as a function of position; ***F***^***a***^ is the local self-propulsion or active force; *D*_*θ*_ is the rotational diffusion coefficient; 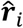 is the direction of particle orientation; and ***ξ*** and ***ξ***_*θ*_ are translational and rotational white noise parameters, respectively.

In our numerical framework, we model the ants as points for which ***v***_*i*_ is always in-line with 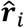. Thus, the translational noise term in **Equation 10** may be taken as zero. Additionally, **Equation 11** may be applied to 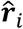 as a random angular change, *ξ*_*i*_ ∈ [**−***πη, π, η*], where *η* is a noise parameter, *η* ∈ [0,1] (*η* = 0 amounts to no random noise, and *η* = 1 results in completely random walks (Shaebani et al., 2020)).

The potential field *U* captures the effects of any external stimuli a surface agent perceives. In the case of ant rafts, we consider this field to be a constant on the bulk where the substrate does not change. Ergo, the only external gradient of *U* occurs at the edges of the raft where the ants perceive water within some detection distance *R*. We will first examine motion on the bulk of the raft.

### Reversion to Vicsek Model on the Bulk

With no gradient of *U* for ants in the center of the raft, the first term inside the parentheses from **Equation 10** goes to zero, giving ***v***_*i*_ = *v*^**−**1^***F***^***a***^. To approximate ***F***^***a***^ we consider our observation that surface ants in direct contact appear to push and entangle one another such that they have some locally shared cooperative force. In our model, we consider a set of *N*_σ_ adjacent surface agents residing within the detection (or contact) distance *R* about the surface agent of interest. If each member of *N*_σ_ propels itself with a force magnitude of *f*^*a*^ in the direction of its own motion (Shaebani et al., 2020), and we roughly assume that these forces are fully transmitted to contact neighbors, then the net propulsion force on a surface agent may be written as 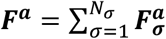, where σ is inclusive of the surface agent of interest. If we take *f*^*a*^ as a constant and 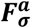 is presumed to act in the direction of motion of each local surface agent, 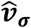, we may rewrite this relation as:

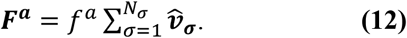

Substituting **Equation 12** into **Equation 10** we get:

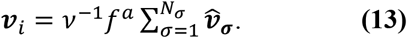

Although this model would theoretically predict unbound speeds for large enough populations of *N*_σ_ moving in unison, we recognize that the physical system is over damped such that the ants have some finite speed due to frictional effects, *v*, which will scale with speed. Furthermore, our framework with 18 DOF naturally constrains the speed of surface agents to the separation distance with its furthest DOF over one time step. Therefore, 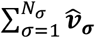 is intrinsically limited by the numerical DOF separation length scale and *f*^*a*^*v*^**−**1^ may be considered a velocity modulus taken as the average ant speed, *v*_0_. This allows us to rewrite **Equation 13** as:

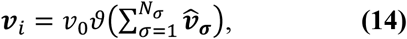

Where ϑ(***V***) denotes normalization of arbitrary vector ***V***. Factoring in random rotation due to rotational noise, *ξ*_*i*_, we may comprehensively rewrite the coupled Langevin equations, **Equation 10** and **Equation 11**, for surface ants on the bulk of the raft as:

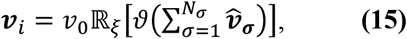

where *ℝ*_*ξ*_[***V***] denotes a random rotation of arbitrary vector ***V*** according to the noise *ξ*_*i*_. **Equation 15** is the form of the Vicsek Model employed by Chaté et al. (2008), and so where ∇_*r*_*U* is zero, we simply employ the Vicsek Model (Vicsek et al., 1995).

For numerical efficacy in our particular framework, we employ the form of the Vicsek Model referenced by Shaebani et al. (2020) of the form:

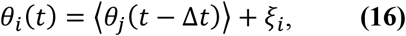

where ⟨*θ*_*j*_(*t* **−** Δ*t*)⟩ is the average angle of the neighboring *N*_σ_ surface agents (including the particle of interest) residing withing distance *R* of the surface agent at time *t* **−** Δ*t*. This form of the Vicsek model was used since the relative angle of each node with respect to each of its neighbors was readily available and easily calculated in our framework. Again, *ξ*_*i*_ is a randomly selected angle within the limits *ξ*_*i*_ ∈ [**−***ηπ, ηπ*], where *η* is our Vicsek Model noise parameter discussed earlier (Shaebani et al., 2020). At the edge of the raft, where **−**∇_*r*_*U* ≠ 0, we must consider not only the active force, but the repulsive edge potential force, as well.

### The Rule of Edge Binding

A surface agent is defined as encountering the edge of the raft when its preferred DOF, as predicted by the Vicsek Model, is a water node. At the edge of the raft, the local change in substrate will result in some effective edge repulsion captured by **−**∇_*r*_*U* in **Equation 10**. To model this edge force, we define some potential between a surface agent and structural agent (*U_ρ_*), and a separate potential between a surface agent and water site (*U*_ω_,). These potentials summarily contribute to **−**∇_*r*_*U*. This edge force is a coarse embodiment of ants’ observed tendency to stay on dry substrates, and may be thought of as a “social force”, as employed in the social force model for pedestrians avoiding external stimuli originally proposed by Helbing and Molnár (1995). Whether or not it is due to surface ants’ affinity to the raft, aversion to water, or both is not immediately clear or relevant, but there is a clear observational tendency for individual ants to avoid moving into the water under their own volition.

Since both the raft and water may influence our edge force, we decompose **−**∇_*r*_*U* as ***F***^*ρ*^ + ***F***^**ω**^, where ***F***^*ρ*^ and ***F***^**ω**^ are the sum of all potential forces exerted on a surface agent by its neighboring raft nodes (*N_ρ_*) and water nodes (*N*_ω_), respectively. These neighbors must be within detection radius *R* about the surface agent of interest. Let us assume that the pair-wise force between a surface agent and the *j*^*th*^ detected neighbor, is a monotonically decreasing potential such that the general form for the total force of that interaction type is 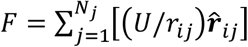, where **r**_*ij*_ is the separation vector between the surface agent and each neighbor. Here, *j* represents either all raft neighbors (the *ρ^th^* neighbor of *N_ρ_*) or water neighbors (the ω^*th*^ neighbor of *N*_ω_). In our numerical framework, wherein *R* is on the order of the contact length scale (*i*.*e*., *R* ∼ 1 𝓁, as determined experimentally) we recognize that our separation distances, *r*_*ij*_, will all be approximately on the order of *R*. Thus, we may rewrite **−**∇_*r*_*U* as:

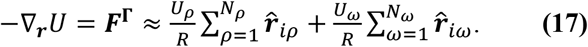

Here, *r*_*iρ*_ and *r*_*i*ω_ are the pair-wise separation vectors between the surface agent and neighboring raft or water nodes, respectively. Since *N_ρ_* and *N*_ω_ are roughly evenly distributed about the center of our circular domain, 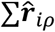 will be roughly equal and opposite to 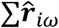, due to symmetry, so **Equation 17** becomes:

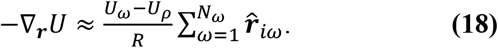

We denote the affinity difference between these two substrates, *U*_ω_ **−** *U_ρ_*, as Γ. Additionally, we denote 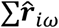 as 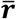 such that **Equation 18** becomes:

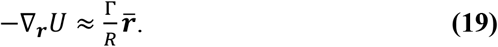

Recall that our propulsion force was provided earlier in **Equation 12**. We multiply this through by *N*_σ_/*N*_σ_ to get 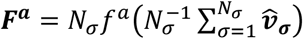, where 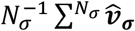 is the local order or cooperativity vector, **φ**_***σ***_, which is **1** when all local particles are moving in the same direction, and → **0** when the local movement is completely disordered (Vicsek and Zafeiris, 2012). Thus, ***F***^***a***^ may be written simply as:

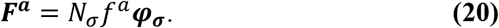

Substituting **Equation 19** and **Equation 20** into **Equation 10** gives the velocity as:

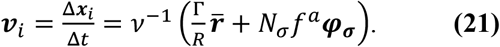

We determine whether the surface agent moves off the raft by projecting **Equation 21** onto the preferred direction of motion (as predicted by the Vicsek Model), 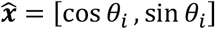, giving:

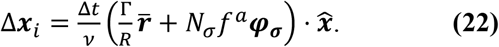

We enforce that if Δ***x***_*i*_ ≤ 0, forwards motion of the surface ant is halted or reversed. We see from **Equation 22** that this occurs when the 1st term is greater than or equal to the 2nd term on the right-hand side. This gives us our fundamental condition for parking, that if 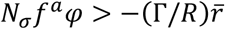, parking occurs where *φ* is the cooperation coefficient or 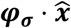 (always positive), and 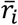 is the edge repulsion alignment coefficient or 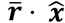⋅(generally negative). Effectively, this states that if the cooperative self-propulsion force in the direction of motion exceeds the edge repulsion force (*i*.*e*., if *F*^*a*^ > **−***F*^Γ^), then edge binding occurs. From this condition, we derive the normalized parking condition:

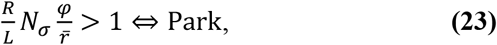

where *L* is the length scale defined by Γ/*f*^*a*^ and physically represents the distance over which a single ant must apply a force *f*^*a*^ to do work equal to the potential difference *U*_ω_ **−** *U*_*r*_. If *U*_ω_ **−** *U*_*r*_ is a constant, *L* is simply inversely proportional to the self-propulsion force. From this length scale, we define the dimensionless activity parameter 𝒜 = 𝓁/*L* = *f*^*a*^𝓁/Γ, which captures the competition between the work an unbound ant does walking its own body length (*f*^*a*^𝓁) and our edge potential (Γ). 𝒜 becomes the free parameter of the model that dictates global expansion and shape change.

### Model Parameters

Our surface agent model has three free parameters (1) *R*, the radius of detection of a single ant (the Vicsek Model radius); (2) 𝒜, the ratio of *f*^*a*^𝓁 to Γ; and (3) *η*, the Vicsek Model noise parameter. *N*_σ_, *φ*, and 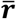 are all measured numerically on a case-by-case basis but are influenced by *R* and *η*. Both *R* and *η* are tuned by comparing our experimental and numerical results using periodic boundary conditions, such that *L* becomes our final sweep parameter. From **Equation 23**, we see that if Γ is assumed constant, increasing 𝒜 effectively requires an increase in the force of ant self-propulsion (*f*^*a*^); thereby, increasing the overall parking rate. In contrast, decreasing 𝒜 will decrease the overall parking rate due to the inverse mechanisms.

### Pausing Surface Traffic

While updating the positions of a surface agent, we run into scenarios where the raft site to which our surface agent would preferentially move deviates greatly from the preferred trajectory predicted by the Vicsek Model. This occurs when all raft sites within the turning limit *θ*_*l*_ are already occupied. In real systems of surface ants, we generally observe that ants whose trajectories are interrupted tend to pause and probe the environment in front of them for on the order of 1 to 10 s before turning around (*i*.*e*., turning greater than *π*/2 radians) to explore elsewhere. Therefore, in cases where no potential raft or water sites meet our set of movement criteria, we do not step the surface agent’s position immediately. Instead, we enforce that the particle pauses for some random time, *t*_*p*_, in the range of [2: 4]Δ*t*. After this time has elapsed, the particle’s orientation is randomly redefined according to the process used at particle initiation. The same is true of surface agents that approach the edge of the raft but are prohibited to edge park because of the limitations invoked through **Equation 23**. These surface agents will also pause for this intermediate timescale before resuming motion in an uncorrelated direction.

The results of this mechanism are to generate traffic jams in confined regions, such as areas with higher local surface densities and those confined by the rafts’ edges (*e*.*g*., the tips of protrusions). Yet this also ensures that these traffic jams (or clusters) are not permanent features in our simulations. Instead, they will dissipate at time scales correlating to both *t*_*p*_ and the cluster or traffic jam size. This practice ensures that clusters occur as they do in experiments yet keeps surface traffic adequately transient.

## Model Calibration

### Pair-wise Strain

Our numerical contractile strain rate was controlled using the parameter, 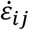, employed according to **Equation 3**. We calibrated 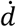 by matching the global decay and numerical unparking rates (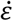 and *δ*, respectively) to those of the experiments. We found that 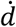 was consistently approximately 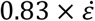, likely due to affine effects within the network structure (Picu, 2011). Additionally, we found that the parking rates matched our experiments when 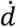 was set according to this observed scaling.

### Vicsek Model Parameters

The effects of altering *R* and *η* are illustrated through the phase table in **Fig. S3**, wherein we show the surface traffic of surface agents’ in our model, occupying a 2D domain with periodic boundary conditions. Moving from left to right, we see that the effect of decreasing *η* (or decreasing the rotational noise, *ξ*_*i*_ in **Equation 16**) is to induce collective motion and directional flow. Likewise, moving from top to bottom, we see that increasing *R* (or increasing the range over which surface agents are influenced by their neighbors) has a similar effect.

**Figure S3:**
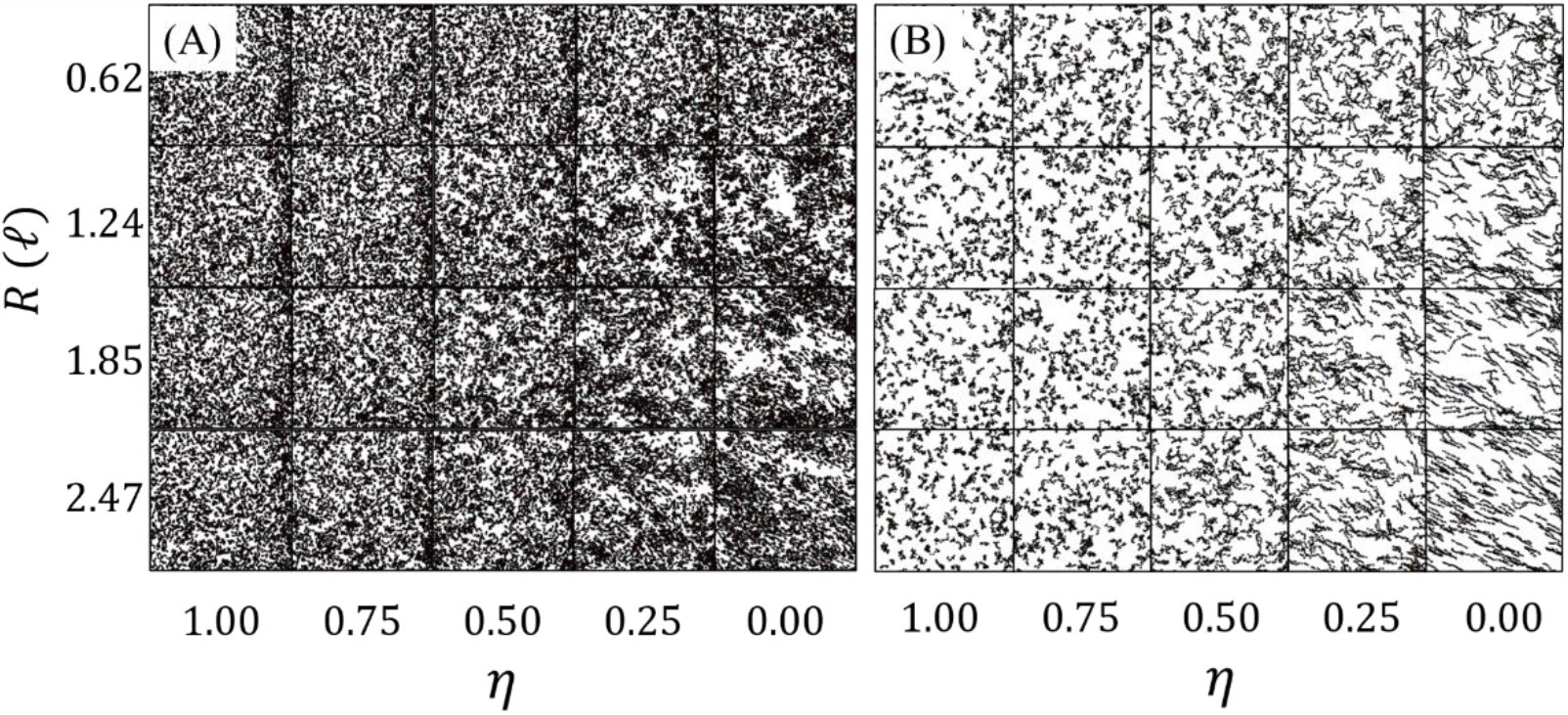
Vicsek Model Phase Diagrams: Snapshots of the surface traffic of surface agents in our numerical model are illustrated at a packing fraction of *ϕ* = 0.239. The full traffic **(A)**, as well as streamlines of just 10% of modeled particles composited from 10 iterations **(B)**, are shown to illustrate the presence of clustering and directional motion, respectively. From top to bottom *R* is swept over the range *R* ∈ [0.62,2.47] 𝓁 and from left to right, *η* is swept over the range *η* = [1.00: **−**0.25: 0].

**Figure S4:**
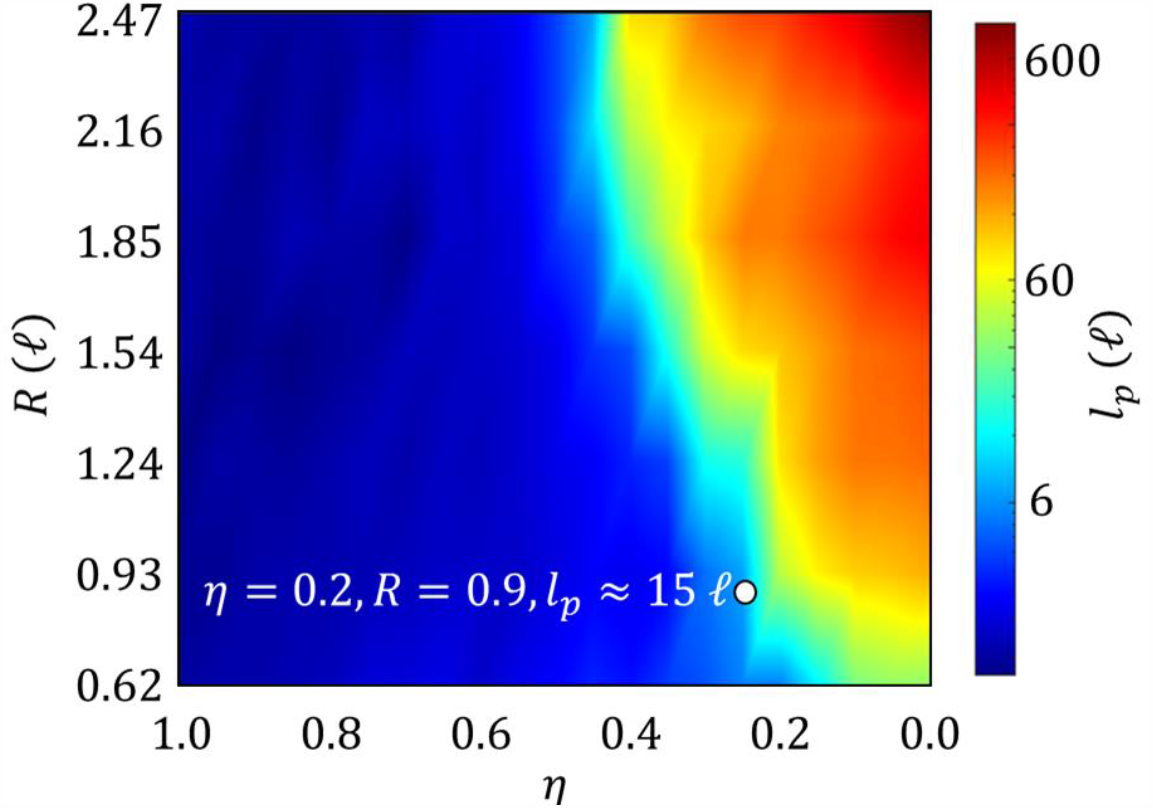
Persistence Length Phase Diagram: An interpolated, 2D heat map illustrates how *l*_*p*_ evolves over the parameter space defined by *R* ∈ [0.62,2.47] 𝓁 and *η* ∈ [0,1] in our numerical model. The point that matches the experimental data is plotted as a white dot.

From our previous work (Wagner et al., 2020), we are aware that *R* is on the order of 1 𝓁, which gives us an estimation of the initial length-scale for our numerical value. Additionally, we found that the surface ant trajectory persistence length, *l*_*p*_, on the bulk of the raft is roughly 15 𝓁. Employing the method used in Wagner et al. (2020), we also calculate *l*_*p*_ for of simulated surface agents in the parameter space given from **Fig. S3**, yielding the heat map illustrated in **Fig. S4**. Matching *R* and *l*_*p*_ to the approximate experimental values of 0.9 𝓁 and 15 𝓁, respectively, we find that the nominal value of *η* ∼ 0.2.

### Parameter Sweep

With *η* set to 0.2, 𝒜 was swept over the range [0.81,3.24]. Although *R* was extrapolated experimentally, it was also swept over the range of [0.62,2.47] 𝓁 to examine the effects of increasing the detection radius on overall raft shape. However, the results presented in the main body of this manuscript hold *R* = 0.9 𝓁 to mimic experimental systems and examine the effects only of local interactions. The extended phase table and heat maps of surface packing fraction, *ϕ*, and peak surface excess, *S*_*max*_, are depicted in **Fig. S5A, S5C** and **S5D**, respectively. **Fig. S5**.**B** depicts the combinations of *R* and 𝒜 that result in matching of *S*_*max*_ (red curve) and *ϕ* (black curve) to those values from the experiments. While the two curves never intersect in the parameter space, this may be attributed to several factors including error in the numerical measurement of surface excess and isotropic detection of neighboring agents within detection radius *R*.

**Figure S5:**
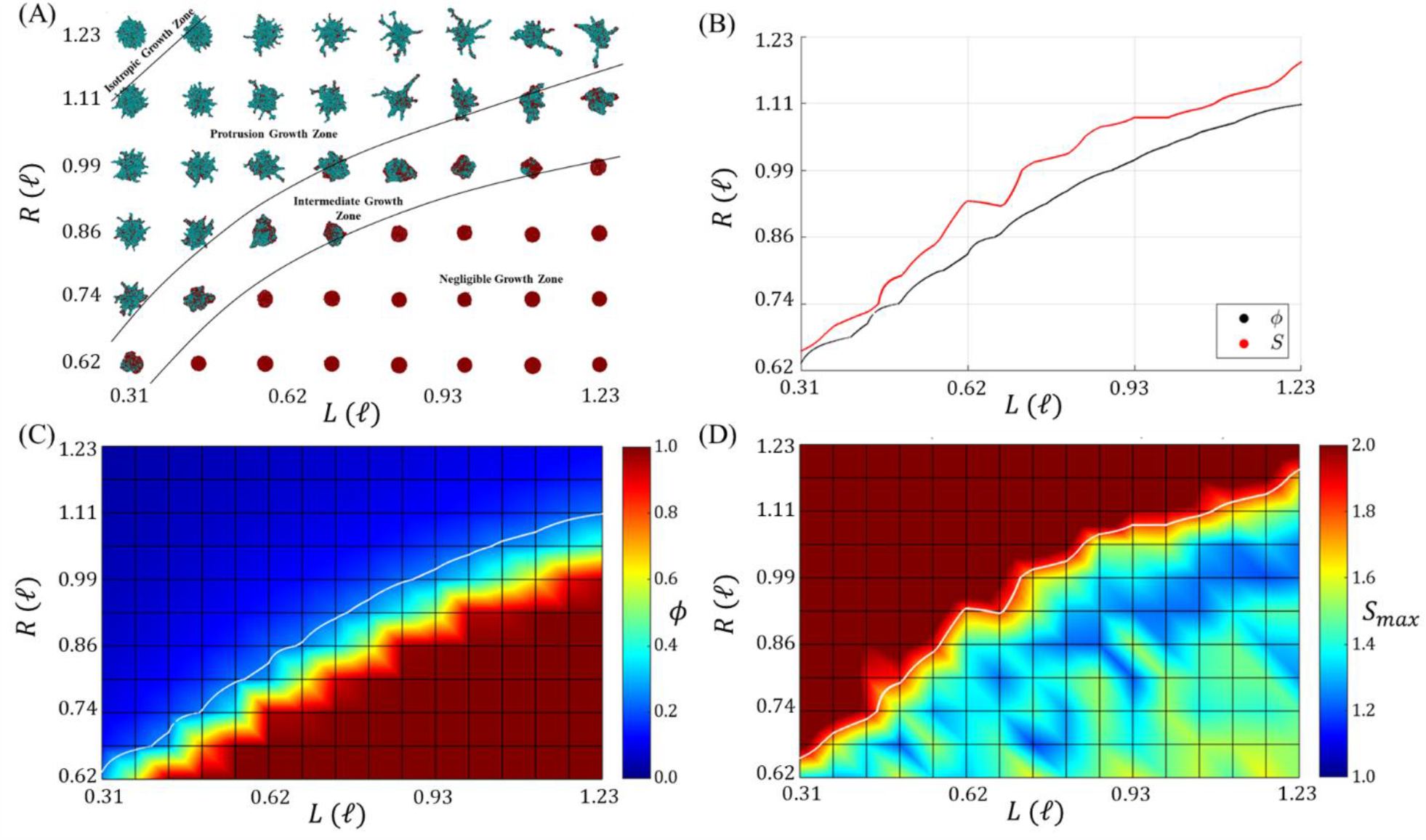
Extended Parameter Sweeps and Phase Diagrams: **(A)** A phase table depicts the morphology of simulated rafts for different values of *R* and 𝒜 after approximately 1 hr. **(B)** Interpolated curves with respect to *R* and 𝒜 depict the phase space in which the maximum surface excess *S*_*max*_ (red) and packing fraction *ϕ* (black) matched those of the experiments (∼ 1.8 and ∼ 0.24, respectively) to within 0.25%. **(C)** A heat map of *ϕ* with respect to *R* and 𝒜 is shown with the white curve corresponding to the black curve from **(B). (D)** A heat map of *S*_*max*_ is shown with respect to *R* and 𝒜 with the white curve corresponding to the red curve from **(B)**.

### Measuring Surface Excess

Recall that surface excess is calculated according to the relation 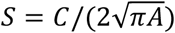, where *C* is the raft’s perimeter length and *A* is the raft’s area. The way in which *C* is measured may significantly impact the estimated value of *S* due to the fractal nature of ant rafts’ edges. Specifically, if the contour length is measured with resolution better than the length scale of the constituents’ size, then *C* will capture the surface roughness of the raft edge and be overestimated in accordance with the coastline paradox (Sokac, 2017). As such, a manual method of measuring *C* by tracing the perimeter of the raft in each frame using ImageJ (Abramoff et al., 2004; Rueden et al., 2017; Schneider et al., 2012) was preferred, as it allowed user discretion in capturing edge defects. This method was used for the experimental dotted line presented in **Fig. 6B**, which coarsely estimates the maximum experimental surface excess on the order of 1.8.

Surface excess of numerical results was estimated by taking *C* as the number of structural agents on the perimeter and *A* as the total number of structural agents. This estimation of *A* is acceptable since our domain density is maintained at 1 node ζ^**−**2^, meaning each structural agent occupies a square space of 1 square unit length. Similarly, this estimation of *C* relies on the fact that the nominal spacing between nodes is ∼ 1 ζ, such that adjacent structural agents in the perimeter may be assumed approximately 1 unit length apart. Perimeter structural agents were defined as agents with neighboring water nodes that reside within the threshold distance of nearest neighbors (*i*.*e*., are ≤ 2 ζ away). This method was used for expediency as it could be automated during simulation post-processing for the ensemble average presented in **Fig. 6B**. However, the assumption of unit spacing (1 ζ) between adjacent edge agents may introduce error in the calculation of *C*.

To directly compare surface excess between experimental and model results, as done in **Fig. 6I**, an alternative and controlled method was used. Both experimental and numerical videos of the raft evolution were imported into ImageJ and converted to binary images with the raft black and the background white. All black pixels that were not part of the raft were removed such that the raft was the only object in the image. Each image was then eroded twelve times to reduce surface roughness at the length scale of individual ants or agents, and then dilated twelve times to revert the rafts back to their original size. The image was then analyzed to measure *A* and *C* and calculate surface excess. Using this method for both the experimental and numerical results allows for a more direct comparison between the two image sources.

It should be noted that regardless of the method employed, the fractal nature of ant rafts’ “coasts” ensures that surface excess is strongly impacted by the resolution with which *C* is measured. While surface excess quantifies shape to some extent, it is best used to interpret qualitative and relative changes in global raft shape rather than draw absolute or quantitative conclusions.

## Data and materials availability

The data supporting the findings of this publication may be reproduced through the agent-based model written in MathWorks® MATLAB R2019b. This model is available at https://doi.org/10.5061/dryad.4f4qrfjb3.

## Acknowledgments

Funding: Franck J. Vernerey gratefully acknowledges the support of the National Science Foundation under Award No. 1761918. The content is solely the responsibility of the authors and does not necessarily represent the official views of the National Science Foundation.

## Supplementary Movie Contents

**Movie S1 – Experimental Treadmilling** presents raw, time-lapsed video footage of an experimental ant raft to visually demonstrate the treadmilling and protrusion growth that takes place. The scale bar represents 10 𝓁 and a timestamp is included in the top right.

**Movies S2-S9** illustrate simulated ant rafts comprised of up to 2,250 agents over a span of 4.5 simulation hours. All rafts were initiated as circles and simulated until pseudo-steady state treadmilling occurred (wherein *α* ≈ *δ*). In all simulations, *R* = 0.9 𝓁 and *η* = 0.2. 𝒜 was set to 1.08, 1.35, 1.42 1.50, 1.59, 1.80, 2.31, and 3.24 for movies **S2-S9**, respectively. Note that a finer separation in 𝒜 between videos is provided in the range 1.35 ≤ 𝒜 ≤ 1.59 since this is the approximate range of continuous phase change identified. All scale bars represent 40 mm or ∼ 14 𝓁.

**Movies S10 – Modulated Activity** illustrates simulated ant raft comprised of 2,250 agents over a span of 5.5 simulation hours when activity was stepped twice between 𝒜 = 1.1 and 𝒜 = 1.6 to demonstrate how activity can toggle a raft between exploratory and inactive phases of protrusion growth and non-growth, respectively. Instantaneous activity level is indicated at the bottom of the video. The raft was initiated as a circle. *R* = 0.9 𝓁 and *η* = 0.2. The scale bar represents 40 mm or ∼ 14 𝓁.

## References

1. Abramoff, M.D., Magalhães, P.J., Ram, S.J., 2004. Image processing with ImageJ [WWW Document]. Biophotonics international. URL http://dspace.library.uu.nl/handle/1874/204900 (accessed 11.17.19).

2. Adams, B.J., Hooper-Bùi, L.M., Strecker, R.M., O’Brien, D.M., 2011. Raft Formation by the Red Imported Fire Ant, Solenopsis invicta. J Insect Sci 11. https://doi.org/10.1673/031.011.17101

3. Alexandre, G., 2015. Chemotaxis Control of Transient Cell Aggregation. J Bacteriol 197, 3230–3237. https://doi.org/10.1128/JB.00121-15

4. Beaune, G., Stirbat, T.V., Khalifat, N., Cochet-Escartin, O., Garcia, S., Gurchenkov, V.V., Murrell, M.P., Dufour, S., Cuvelier, D., Brochard-Wyart, F., 2014. How cells flow in the spreading of cellular aggregates. Proc Natl Acad Sci U S A 111, 8055–8060. https://doi.org/10.1073/pnas.1323788111

5. Bugyi, B., Carlier, M.-F., 2010. Control of actin filament treadmilling in cell motility. Annu Rev Biophys 39, 449–470. https://doi.org/10.1146/annurev-biophys-051309-103849

6. Chaté, H., Ginelli, F., Grégoire, G., Peruani, F., Raynaud, F., 2008. Modeling collective motion?: variations on the Vicsek model.

7. Cini, R., Loglio, G., Ficalbi, A., 1972. Temperature dependence of the surface tension of water by the equilibrium ring method. Journal of Colloid and Interface Science 41, 287–297. https://doi.org/10.1016/0021-9797(72)90113-0

8. De Magistris, G., Marenduzzo, D., 2015. An introduction to the physics of active matter. Physica A: Statistical Mechanics and its Applications, Proceedings of the 13th International Summer School on Fundamental Problems in Statistical Physics 418, 65–77. https://doi.org/10.1016/j.physa.2014.06.061

9. Deblais, A., Woutersen, S., Bonn, D., 2020. Rheology of Entangled Active Polymer-Like T. Tubifex Worms. Phys. Rev. Lett. 124, 188002. https://doi.org/10.1103/PhysRevLett.124.188002

10. Dong, C., Aznavoorian, S., Liotta, L.A., 1994. Two phases of pseudopod protrusion in tumor cells revealed by a micropipette. Microvasc. Res. 47, 55–67. https://doi.org/10.1006/mvre.1994.1005

11. DuChez, B.J., Doyle, A.D., Dimitriadis, E.K., Yamada, K.M., 2019. Durotaxis by Human Cancer Cells. Biophysical Journal 116, 670–683. https://doi.org/10.1016/j.bpj.2019.01.009

12. Elgeti, J., Gompper, G., 2013. Wall accumulation of self-propelled spheres. EPL 101, 48003. https://doi.org/10.1209/0295-5075/101/48003

13. Fily, Y., Baskaran, A., Hagan, M.F., 2014. Dynamics of self-propelled particles under strong confinement. Soft Matter 10, 5609–5617. https://doi.org/10.1039/C4SM00975D

14. Helbing, D., Molnár, P., 1995. Social force model for pedestrian dynamics. Phys. Rev. E 51, 4282–4286. https://doi.org/10.1103/PhysRevE.51.4282

15. Hu, D.L., Phonekeo, S., Altshuler, E., Brochard-Wyart, F., 2016. Entangled active matter: From cells to ants. Eur. Phys. J. Spec. Top. 225, 629–649. https://doi.org/10.1140/epjst/e2015-50264-4

16. Loi, D., Mossa, S., Cugliandolo, L.F., 2008. Effective temperature of active matter. Phys. Rev. E 77, 051111. https://doi.org/10.1103/PhysRevE.77.051111

17. Méhes, E., Vicsek, T., 2014. Collective motion of cells: from experiments to models. Integr. Biol. 6, 831–854. https://doi.org/10.1039/C4IB00115J

18. Mlot, N.J., Tovey, C.A., Hu, D.L., 2011. Fire ants self-assemble into waterproof rafts to survive floods. PNAS 108, 7669–7673. https://doi.org/10.1073/pnas.1016658108

19. Neuhaus, J.-M., Wanger, M., Keiser, T., Wegner, A., 1983. Treadmilling of actin. J Muscle Res Cell Motil 4, 507–527. https://doi.org/10.1007/BF00712112

20. Nikola, N., Solon, A.P., Kafri, Y., Kardar, M., Tailleur, J., Voituriez, R., 2016. Active Particles with Soft and Curved Walls: Equation of State, Ratchets, and Instabilities. Phys. Rev. Lett. 117, 098001. https://doi.org/10.1103/PhysRevLett.117.098001

21. Peleg, O., Peters, J.M., Salcedo, M.K., Mahadevan, L., 2018. Collective mechanical adaptation of honeybee swarms. Nature Phys 14, 1193–1198. https://doi.org/10.1038/s41567-018-0262-1

22. Picu, R.C., 2011. Mechanics of random fiber networks - A review. Soft Matter 7, 6768–6785. https://doi.org/10.1039/c1sm05022b

23. Poujade, M., Grasland-Mongrain, E., Hertzog, A., Jouanneau, J., Chavrier, P., Ladoux, B., Buguin, A., Silberzan, P., 2007. Collective migration of an epithelial monolayer in response to a model wound. Proc. Natl. Acad. Sci. U.S.A. 104, 15988–15993. https://doi.org/10.1073/pnas.0705062104

24. Reichhardt, C., Thibault, J., Papanikolaou, S., Reichhardt, C.J.O., 2018. Laning and clustering transitions in driven binary active matter systems. Phys. Rev. E 98, 022603. https://doi.org/10.1103/PhysRevE.98.022603

25. Rueden, C.T., Schindelin, J., Hiner, M.C., DeZonia, B.E., Walter, A.E., Arena, E.T., Eliceiri, K.W., 2017. ImageJ2: ImageJ for the next generation of scientific image data. BMC Bioinformatics 18, 529. https://doi.org/10.1186/s12859-017-1934-z

26. Ryan, G.L., Watanabe, N., Vavylonis, D., 2012. A review of models of fluctuating protrusion and retraction patterns at the leading edge of motile cells. Cytoskeleton 69, 195–206. https://doi.org/10.1002/cm.21017

27. Schneider, C.A., Rasband, W.S., Eliceiri, K.W., 2012. NIH Image to ImageJ: 25 years of image analysis. Nature Methods 9, 671–675. https://doi.org/10.1038/nmeth.2089

28. Shaebani, M.R., Wysocki, A., Winkler, R.G., Gompper, G., Rieger, H., 2020. Computational models for active matter. Nature Reviews Physics 2, 181–199. https://doi.org/10.1038/s42254-020-0152-1

29. Simon, C., Kusters, R., Caorsi, V., Allard, A., Abou-Ghali, M., Manzi, J., Di Cicco, A., Lévy, D., Lenz, M., Joanny, J.-F., Campillo, C., Plastino, J., Sens, P., Sykes, C., 2019. Actin dynamics drive cell-like membrane deformation. Nat. Phys. 15, 602–609. https://doi.org/10.1038/s41567-019-0464-1

30. Sokac, A.M., 2017. Seeing a Coastline Paradox in Membrane Reservoirs. Developmental Cell 43, 541–542. https://doi.org/10.1016/j.devcel.2017.11.013

31. Tennenbaum, M., Fernandez-Nieves, A., 2020. Activity effects on the nonlinear mechanical properties of fire-ant aggregations. Phys. Rev. E 102, 012602. https://doi.org/10.1103/PhysRevE.102.012602

32. Trepat, X., Chen, Z., Jacobson, K., 2012. Cell Migration. Compr Physiol 2, 2369–2392. https://doi.org/10.1002/cphy.c110012

33. Vicsek, T., Czirók, A., Ben-Jacob, E., Cohen, I., Shochet, O., 1995. Novel Type of Phase Transition in a System of Self-Driven Particles. Physical Review Letters 75, 1226.

34. Vicsek, T., Zafeiris, A., 2012. Collective motion. Physics Reports 517, 71–140. https://doi.org/10.1016/j.physrep.2012.03.004

35. Vutukuri, H.R., Hoore, M., Abaurrea-Velasco, C., van Buren, L., Dutto, A., Auth, T., Fedosov, D.A., Gompper, G., Vermant, J., 2020. Active particles induce large shape deformations in giant lipid vesicles. Nature 586, 52– 56. https://doi.org/10.1038/s41586-020-2730-x

36. Wagner, R.J., Such, K., Hobbs, E., Vernerey, F.J., 2020. Collective treadmilling in fire ant rafts permits sustained protrusion growth. arXiv:2012.15782 [cond-mat, physics:physics].

37. Wen, J.H., Choi, O., Taylor-Weiner, H., Fuhrmann, A., Karpiak, J.V., Almutairi, A., Engler, A.J., 2015. Haptotaxis is cell type specific and limited by substrate adhesiveness. Cell Mol Bioeng 8, 530–542. https://doi.org/10.1007/s12195-015-0398-3

38. Zimmerman, S.P., Asokan, S.B., Kuhlman, B., Bear, J.E., 2017. Cells lay their own tracks – optogenetic Cdc42 activation stimulates fibronectin deposition supporting directed migration. J Cell Sci 130, 2971–2983. https://doi.org/10.1242/jcs.205948

